# Prospective evaluation of structure-based simulations reveal their ability to predict the impact of kinase mutations on inhibitor binding

**DOI:** 10.1101/2024.11.15.623861

**Authors:** Sukrit Singh, Vytautas Gapsys, Matteo Aldeghi, David Schaller, Aziz M. Rangwala, Jessica B. White, Joseph P. Bluck, Jenke Scheen, William G. Glass, Jiaye Guo, Sikander Hayat, Bert L. de Groot, Andrea Volkamer, Clara D. Christ, Markus A. Seeliger, John D. Chodera

## Abstract

Small molecule kinase inhibitors are critical in the modern treatment of cancers, evidenced by the existence of over 80 FDA-approved small-molecule kinase inhibitors. Unfortunately, intrinsic or acquired resistance, often causing therapy discontinuation, is frequently caused by mutations in the kinase therapeutic target. The advent of clinical tumor sequencing has opened additional opportunities for precision oncology to improve patient outcomes by pairing optimal therapies with tumor mutation profiles. However, modern precision oncology efforts are hindered by lack of sufficient biochemical or clinical evidence to classify each mutation as resistant or sensitive to existing inhibitors. Structure-based methods show promising accuracy in retrospective benchmarks at predicting whether a kinase mutation will perturb inhibitor binding, but comparisons are made by pooling disparate experimental measurements across different conditions. We present the first prospective benchmark of structure-based approaches on a blinded dataset of in-cell kinase inhibitor affinities to Abl kinase mutants using a NanoBRET reporter assay. We compare NanoBRET results to structure-based methods and their ability to estimate the impact of mutations on inhibitor binding (measured as ΔΔG). Comparing physics-based simulations, Rosetta, and previous machine learning models, we find that structure-based methods accurately classify kinase mutations as inhibitor-resistant or inhibitor-sensitizing, and each approach has a similar degree of accuracy. We show that physics-based simulations are best suited to estimate ΔΔG of mutations that are distal to the kinase active site. To probe modes of failure, we retrospectively investigate two clinically significant mutations poorly predicted by our methods, T315A and L298F, and find that starting configurations and protonation states significantly alter the accuracy of our predictions. Our experimental and computational measurements provide a benchmark for estimating the impact of mutations on inhibitor binding affinity for future methods and structure-based models. These structure-based methods have potential utility in identifying optimal therapies for tumor-specific mutations, predicting resistance mutations in the absence of clinical data, and identifying potential sensitizing mutations to established inhibitors.

## INTRODUCTION

The emergence of drug-resistant mutations represents a significant obstacle in the effective treatment of various diseases, including cancer. This is most recently exemplified by over 80 FDA-approved kinase inhibitors (also known as TKIs) already having known resistant mutations.^1^ These mutations can arise in the target proteins of therapeutic agents, rendering them less susceptible or completely resistant to the drugs’ intended mechanisms of action. The ability to predict drug-resistant mutations in advance promises to guide precision medicine.^2–4^

A number of susceptible and resistance mutations have been characterized for FDA-approved kinase inhibitors (e.g. OncoKB) but a much larger number of mutations in the target of therapy (the kinase domain) have been observed for which no data is available.^4–9^

Predicting the impact of mutations remains challenging due to the complex and diverse nature of resistance mechanisms. Kinase mutations may decrease drug-binding affinity or potency,^10–12^ increase kinase activity,^13–16^ tune inhibitor sensitivity profiles,^11,17,18^ or any combination of these mechanisms, or other mechanisms involving additional cellular machinery.^19,20^ Alternatively, mutations may shift the population of conformations towards alternative states, decreasing the population of drug-compatible conformations.^21–26^ Some resistance mutations may even be compensatory, increasing activity of the kinase either through shifting towards increased propensity towards an active kinase,^27,28^ or increasing the affinity for ATP.^14,22,29^ Of these potential mechanisms, the most direct way that a mutation can cause drug resistance is by perturbing the kinase-inhibitor binding affinity.^10–12,30,31^

Mutation-induced changes in kinase-inhibitor binding affinity would lead to reduced drug potency. In some cases, a change in binding affinity upon mutation may indicate that a mutation *sensitizes* a protein to an inhibitor, offering new therapeutic strategies.^17,32,33^ Thus, being able to measure the impact of a mutation on protein-inhibitor binding affinity offers new insights and avenues into circumventing drug resistance.

Experimental methods such as mutagenesis studies and binding assays can provide critical biophysical insight but are often time-consuming, costly, and limited in throughput.^28,34,35^ One approach to mechanistically characterizing drug-resistant mutations involves measuring the direct change in binding free energy (ΔΔG).^10–12,31,36^

Structure-informed methods show promise in predicting the impact of mutations on ΔΔG, and may help classify mutants as resistant or susceptible to kinase inhibitors. Existing benchmarks have only been retrospective.^10,12,31^ Computational approaches have been previously shown to predict changes in binding affinity caused by mutations with an average error of 1.2 to 2.0 kcal/mol.^10,37^ Rosetta-based methods predict ΔΔG using Monte Carlo methods to sample and optimize protein conformations, predicting mutational effects on protein structure and stability.^10,12,38^ However Monte Carlo search methods are limited in their ability to sample conformational changes upon mutation and so may not capture impact of a mutation on the conformational landscape of a protein. Molecular dynamics (MD)-based use atomistic detail to describe the dynamics and energetics of protein-drug complexes.^23,39,40^ By simulating the behavior of the system over time, molecular dynamics can capture the effects of mutations on the stability and dynamics of the binding pocket and the interactions with the drug.^23,33,41^ However these methods require extensive simulation to appropriately sample larger clinically-relevant systems, limiting throughput.^39^ Recently, machine learning (ML) methods have also gained prominence in predicting ΔΔG changes in binding affinity.^12,31,42^ Such ML approaches utilize large datasets of experimentally determined binding affinities to train predictive models.^10,12^ By learning the relationship between sequence, structure, and binding affinity, ML models can predict the impact of mutations on binding affinity with remarkable accuracy. However, these methods require large datasets of clinically identified mutations, and knowledge of their mechanistic impact to accurately predict mutational impact.^43,44^ As such, ML methods require an existing dataset to train upon using consistent experimental measurements, which does not exist yet. Once such datasets become available, active learning loops can be deployed to cycle between experimental measurement, model training, and predictive evaluation to improve ML model prediction further.^45^

Alchemical methods, sometimes referred more generally to as Free Energy Calculations (FECs) in prior work (Fig. S1),^46^ have emerged as powerful computational methods for predicting changes in binding affinity without the need for extensive experimental data that requires a large amount of time and cost to generate.^31,36,37,39,46,47^ Such structure-based alchemical methods aim to complement existing experimental measurements via large-scale parallel assessment of many mutations prior to experimental measurement.

These alchemical methods are used to estimate free energies of binding (ΔG) by estimating the energetic cost of atoms going from one thermodynamic configuration to another via tuning the strength of atomic interactions via a so-called alchemical transformation.^46^ To study the impact of a mutation on drug binding, we estimate the energetic cost of transforming amino acids between wild-type (WT) and mutant residues in the presence and absence of a ligand (Fig. S1). These alchemical methods compute free energy differences (ΔG) using a so-called “transformation” between the wild-type and mutant amino acid atoms. This transformation is “alchemical” in that it scales, using the parameter λ, bonded and nonbonded interactions of an amino acid sidechain found in WT and mutant. Briefly, alchemical free energy methods estimate the impact of a mutation on drug binding by estimating the thermodynamic cost of transforming one amino acid to another in the presence and absence of a ligand. These alchemical methods compute free energy differences (ΔG) using a so-called “transformation” between the wild-type and mutant amino acid atoms. This transformation is “alchemical” in that it scales, using the parameter λ, bonded and nonbonded interactions of an amino acid sidechain found in WT and mutant Abl kinase (Fig. 3) such that λ=0 represents a complete WT Abl and λ=1 represents the mutant; values of 0<λ<1 are a scaled combination of the two sets of interactions. During a transformation, the sidechain of one amino acid (WT) is “phased out” while another amino acid chain (the mutant) is “phased in” (Fig. 3).^37,46–48^ In computing the free energy (ΔG) in both holo and apo conditions, a thermodynamic cycle is constructed, in which we subtract the apo ΔG from the holo ΔG to arrive at the thermodynamic impact of mutation upon drug-binding (ΔΔG) (Fig. S1).^46^ Several alchemical approaches can be used to estimate ΔΔG changes associated with mutations. Replica exchange methods, such as Hamiltonian replica exchange molecular dynamics (RepEx, HREX, or REMD),^47,49–52^ enable enhanced sampling of conformational space and provide insights into the thermodynamics of the binding process. Non-equilibrium switching methods utilize fast out-of-equilibrium transitions along the reaction coordinate to estimate free energy differences.^36,53,53–56^ However, existing benchmarks have only been retrospective,^10,12,36,56,57^ and the prospective capabilities of FECs require examination.

Previous FEC benchmarks derive from IC50 measurements across multiple types of biochemical or in cellular measurements,^18,24^ ranging from qPCR detection, Ba/F3 activity assays, or mobility shift assays (gel shift).^30,58–61^ In turn, there are many more sources of experimental variation that can affect the quality of benchmarking measurements.^62^ Furthermore, variation in experimental contexts, from cells to in vitro mobility measurements, to variability in cell type for qPCR, may cause context-dependent variation that alters the measured impact of clinical kinase mutations.^62,63^ Such protocol specific variability may not capture accurate cancer-like cellular contexts and environments, thus introducing noise into the benchmarking dataset.^64,65^ Ideally, our ability to measure IC50 would use a consistent experimental readout that more directly and consistently reports on binding. There is a need for comparing against a dataset that uses a single type of measurement in a controlled rigorous manner.

NanoBRET offers a high throughput in-cell approach to prospectively evaluate mutations and their impact on kinase-inhibitor binding.^66,67^ This method relies on the principle of resonance energy transfer between a bioluminescent donor and a fluorescent acceptor. NanoBRET has proven valuable for high-throughput screening of mutations and their effects on ligand binding,^24,68,69^ and allows for the cellular environment to influence kinase-inhibitor binding, providing an in-cell characterization of the change in affinity.^24,68^ The consistency in experimental measurements allows us to avoid multiple sources of experimental variation when benchmarking FEC results.^63^ NanoBRET data has previously demonstrated its capacity to measure the impact of mutations on Abl kinase, even revealing new possible kinetic mechanisms of drug resistance.^24^ Computational tools to identify these highly sensitive mutations *a priori* may reduce patient burden and increase treatment outcomes as we are able to more precisely treat patients who would benefit most from the use of sensitized drugs.

Here, we present the first prospective benchmark of structure-informed physical and machine learning models for the prediction of resistance/susceptibility, using a NanoBRET assay to measure the impact of clinical Abl mutations within cells, considering first- and second-generation TKIs, imatinib and dasatinib (Fig. 1 and 2).^24,61,70–72^ We compare and evaluate the performance of structure-based methods to predict the impact of mutations, including replica exchange and non-equilibrium switching methods, Rosetta-based methods,^38^ and a previously published Random Forest method (Fig. 3).^10,57^ The Rosetta flex_ddg protocol models the structural and energetic effects of mutations by generating mutant structures and extensively sampling possible rotamers. The protocol then iteratively refines ΔΔG across multiple iterations using the Rosetta scoring function to generate averaged predictions that account for likely rotamers of both WT and mutant amino acid.^38^ A Random Forest model uses an ensemble of randomized decision trees to predict the change in binding affinity (ΔΔG) of drugs upon point mutations in the human Abl kinase.^10,12,73^ The model was trained on a curated dataset of 144 ΔΔG values for eight tyrosine kinase inhibitors, incorporating a diverse set of features that describe ligand properties, mutation environments, amino acid changes, protein-ligand interactions, docking scores, and solvent accessibility.^31,58,60,61^

**Figure 1.**
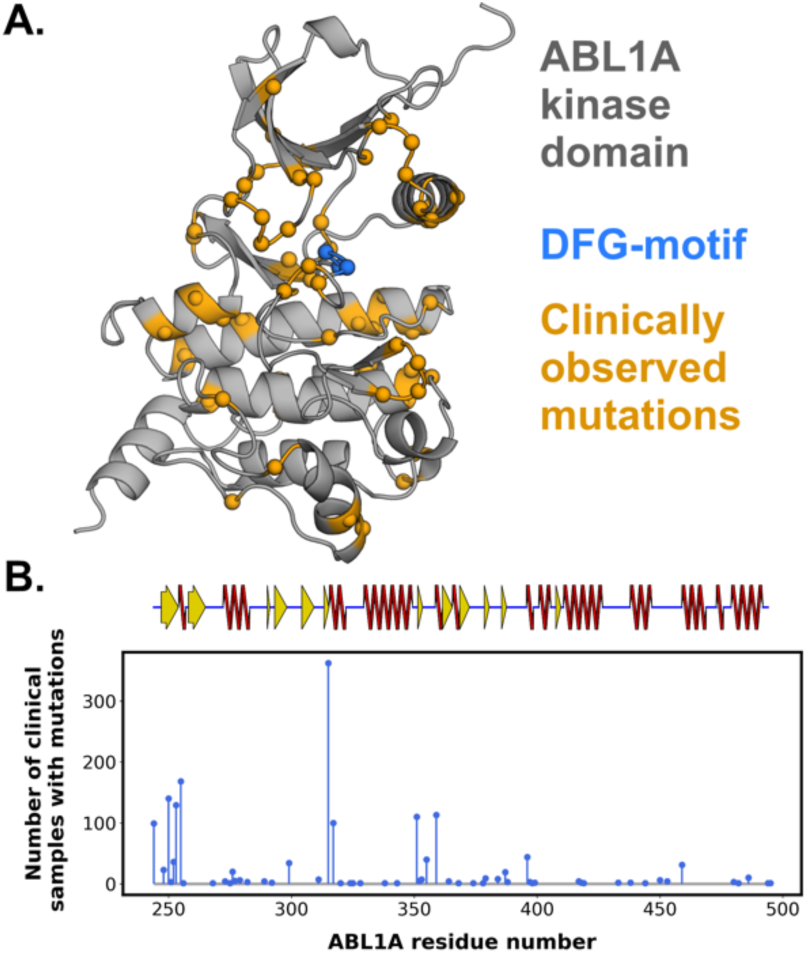
Clinically-relevant mutations occur throughout Abl kinases, at varying frequencies. **A.** Structure of Abl kinase (PDB: 1OPJ), highlighting the positions of clinically-derived Abl mutations (orange spheres) associated with resistance (defined by COSMIC or OncoKB). **B.** Lollipop plot denoting the number of clinical samples found in the Cosmic/cBioPortal Databases denoting the number of times a mutation occurs at each position of the kinase domain sequence. Secondary structure of the corresponding region is indicated (above) denoting whether the region is helical (red), a β-strand (yellow) or a loop (blue).

**Figure 2:**
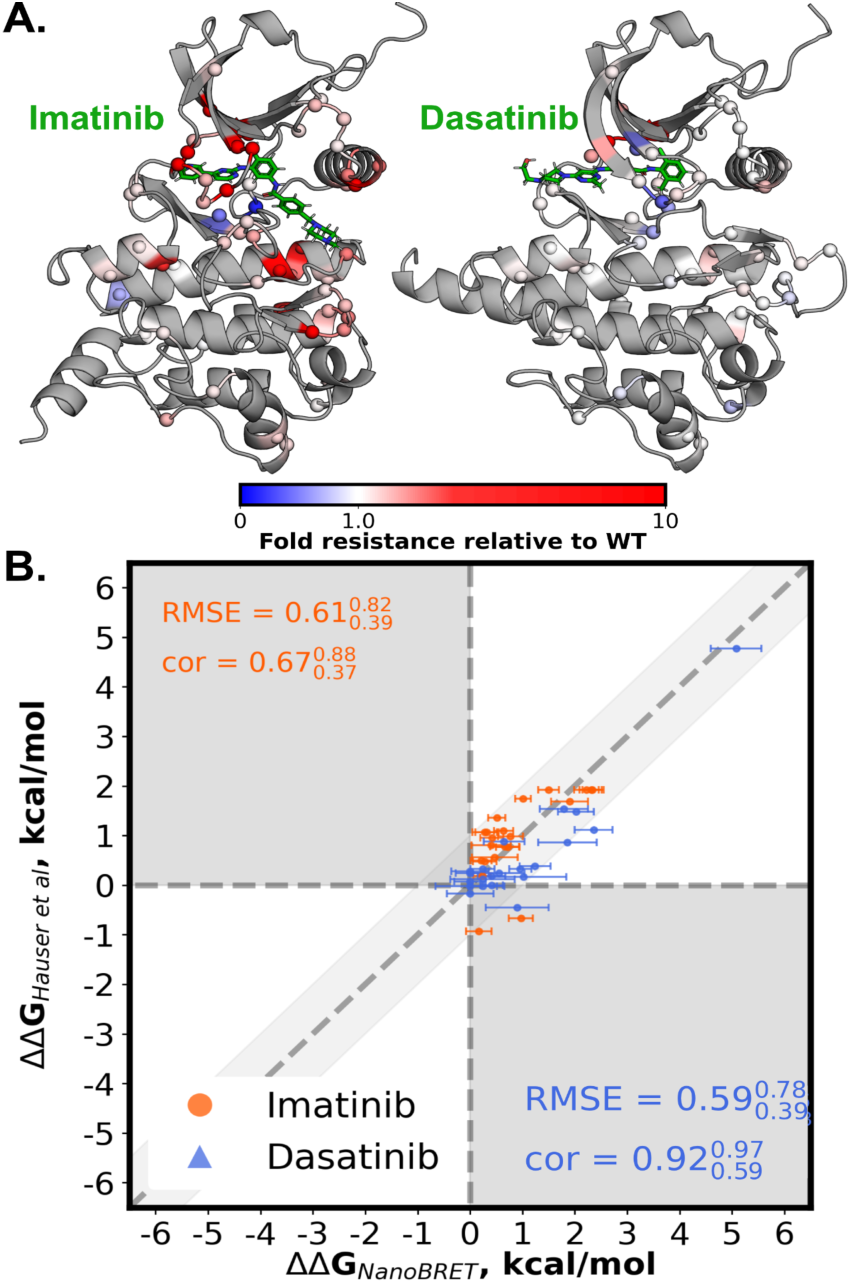
An intracellular NanoBRET assay for measuring the impact of clinical mutations on kinase inhibitor binding free energy is concordant with previously-reported IC50-derived data. **A.** Experimental IC50 values of Abl kinase binding to Imatinib (left, green) or Dasatinib (right, green) for a variety of mutations (spheres) normalized by the WT IC50 of each compound (color scale, bottom). This WT-based normalization demonstrates which mutations are resistant or sensitizing to either imatinib or dasatinib. **B.** Experimental ΔΔG values from the current work plotted against those from Hauser et al.^31^ A subset of 18 mutations common to the sets of Hauser et al.^31^ and the current work is considered. Data points for which the experimental values disagree by more than 1 kcal/mol are marked in a darker edge color. Root Mean Square Error (RMSE) and Pearson correlation (labeled “cor”) between datasets noted for each mutation-ligand pair, alongside 95% confidence intervals.

In this work, we apply these methods on clinically identified mutations of Abl kinase, and compare ΔΔG predictions against measured values from experimental nanoBRET measurements of Abl binding to imatinib and dasatinib. We show that high throughput biophysical methods like nanoBRET provide comparable measurements to other low-throughput methods, demonstrating the value of nanoBRET as a benchmark measurement for assessing ΔΔG accuracy. We prospectively evaluate 90 clinically observed mutations of Abl kinase, evaluating the ability of computational methods to predict ΔΔG prospectively. We also retrospectively evaluate modes of failure behind two clinically occurring, difficult-to-predict mutations using open-source tools that can be massively parallelized for ease of study. We demonstrate the capacity of free energy calculations to independently predict accurate ΔΔG values even for distal mutations that are not near the binding site.

Lastly, we also demonstrate that FECs are capable of improving ΔΔG by considering alternative protonation states. The integration of multiple computational methods allows for a comprehensive and reliable prediction of mutation effects, aiding in the design of effective therapeutic strategies and advancing precision medicine.

## METHODS

### Lollipop plot generation of clinical variants

All variants were obtained from the Catalogue of Somatic Mutations in Cancer (COSMIC) ABL1 entry (ENST00000318560) as of 7/28/2023.^5^ Counts were aggregated for the 94 variants assessed in this study.

### Prospective Free energy calculations

#### Free energy calculations with FEP+

System preparation and free energy calculations were performed using Schrödinger Maestro Suite Version 2020-3 (Schrödinger Release 2020-3: Maestro, Prime, FEP+; Schrödinger, LLC: New York, 2020).^85^ System preparation used the protein preparation wizard within Maestro and resembled Hauser et. al. but differed in some aspects which will be pointed out below.^31^ Residue numbering was adjusted such that the threonine gatekeeper of Abl kinase obtained the residue number 315 to match common practice. As in Hauser et. al. chain B of PDB structure 1OPJ was used as input for the calculations of imatinib in complex with Abl.^72^ However, the complete chain as present in the PDB file was used. Solvent exposed serine 336 (355 in PDB file) was mutated to asparagine using Maestro. Termini were capped and all water molecules present in the PDB structure were kept. No loops were missing and needed modeling.

Imatinib was modeled as positively charged by the protein preparation wizard, labeled as the “imatinib+1” species. For calculations investigating the impact of inhibitor charge, imatinib was manually neutralized to generate the “imatinib+0” species. For the simulations of dasatinib in complex with Abl PDB structure 4XEY was used.^103^ Specifically, Chain A of the PDB structure (4XEY) was used, but no homology modelling was done to add on unresolved N- and C-terminal residues.

Both imatinib and dasatinib ligand parameters are described by the OPLS forcefield.^104^ The preparation wizard neutralized aspartates 381 and 421. Input structures for imatinib and dasatinib can be found as a part of the supplementary material and are available online (https://osf.io/s6ktq/). The effect of residue mutation on ligand affinity was calculated using FEP+ with default settings i.e. using the muVT ensemble, a simulation time of 5ns per lambda window and 12, 16, or 24 lambda windows for standard, core hopping, and charge changing perturbations, respectively. Each run was performed in triplicates using different random number seeds. Error estimates correspond to standard deviations over the three runs.

#### Curation and preparation of structures for Nonequilibrium protocols with PMX (NEQ)

Structures of the ABL1A-inhibitor complexes were used as previously described by drawing up on previously published crystal structures (4XEY and 1OPJ for dasatinib and imatinib, respectively).^31^ Apo structures were generated by discarding ligand atoms, and crystallographic water molecules were retained. All mutant structures were generated using FoldX v4.^105^ Amino acid protonation states were set at pH 7.4 using PDB2PQR and PROPKA v3.1 via the HTMD protein preparation tool (v1.12).^106–109^ Ligand protonation states were kept previously described in Hauser et al by default.^31^ Proteins were described using the Amber99sb*-ILDN (shown as “A99”) and Amber14sb (shown as “A14sb”) force fields.^110–113^ The TIP3P water model was used.^114^ Ligands parameters were created using GAFF2 (v2.1) via AmberTools 16^115^ where charges were described using restrained electrostatic potential (RESP).^116^ Gaussian 09 (Rev D.01) was used to conduct geometry optimizations and molecular electrostatic potential (ESP) calculations using the HF/6-31G* level of theory.

Three optimization steps were used to ensure the ligand conformation remained similar to the kinase-bound poses. ESP points were sampled according to the Merz-Kollman scheme.^117,118^ Halogen atom σ-holes were modeled as previously described by Kolář and Hobza.^119^ Specifically, we add off-site charge modifications to the GAFF force field that alter halogen atoms to more accurately model σ-hole interactions between protein amino acids and halogens. This modification is done to appropriately model halogen atoms in the small molecule inhibitors in this study. All ligand parameters can be found in the input files in (https://osf.io/s6ktq/). Protein-ligand systems were solvated in a dodecahedral box with periodic boundary conditions with a minimum padding distance 12 Å from the protein system to the edge of the box. Sodium and chloride ions were added to neutralize the wild-type (WT) system at a concentration of 0.15 M NaCl. For the mutants, the same number of ions as in the wild type systems was added; i.e. the net charge of the wild type systems was always zero, while the net charge of the mutant systems was allowed to deviate from zero. Clashes may also be present in this initial structure as FoldX does not consider the presence of ligands when inserting mutations in the protein.

Clashes were considered present if any protein heavy atom was within 1.5 Å of any ligand heavy atom. If one or more clashes were present, an approach similar to alchembed was used to resolve them: 2000 steepest descent minimization steps were conducted, after which, the ligand vdW interactions were switched on for 2000 additional steps in using the MD integrator steps carried out with a 0.5 femtosecond time step with position restraints (at 1000 kJ mol^−1^ nm^−2^) on all heavy atoms.^120^

#### Non-equilibrium protocol (NEQ) with PMX

All simulations were carried out on Gromacs 2016 on Intel Xeon processors with Ivy Bridge (4 cores, E3-1270 v2) or Broadwell (10 cores, E5-2630 v4) architectures and NVIDIA GeForce GPUs (GTX 1070, GTX 1080, or GTX 1080 Ti).^121,122^ Energy minimization was carried out using a steepest descent algorithm for 10,000 steps. The systems were subsequently simulated for 100 ps in the isothermal-isobaric ensemble (NPT) with harmonic position restraints applied to all solute heavy atoms with a force constant of 1000 kJ mol^-1^ nm^-2^. Equations of motion were integrated with a leap-frog integrator and a time-step of 2 femtoseconds (fs). The temperature was coupled with the stochastic v-rescale thermostat at the target temperature of 300 K.^123^ The pressure was controlled with the Berendsen weak coupling algorithm at a target pressure of 1 bar.^124^ The particle mesh Ewald (PME) algorithm was used for electrostatic interactions with a real space cut-off of 10 Å when using Amber force fields, a spline order of 4, a relative tolerance of 10^−5^, and a Fourier spacing of 1.2 Å.^125^ Verlet cut-off schemes with the potential-shift modifier was used with a Lennard-Jones interaction cut-off of 10 Å, and a buffer tolerance of 0.005 kJ mol^−1^ ps^−1^.^126^ All bonds were constrained with the P-LINCS algorithm.^127^ For equilibration, unrestrained MD simulations were then performed for 1ns in the NPT ensemble with the Parrinello-Rahman barostat at 1 bar with a time constant of 2 ps.^128^ Production simulations were then performed for 3 ns for A14 and 5 ns for A99.

For ΔΔG estimations, the above procedure for equilibrium simulations was repeated ten times on both the apo and complex states of both wild-type and mutant to estimate ΔΔG. From each of these ten equilibrium simulations, 30 equally spaced frames were extracted to serve as starting configurations for the non-equilibrium protocol, generating a total of 300 non-equilibrium trajectories. There were 150 trajectories going from wild-type to mutant (“forward”) and 150 trajectories going from mutant to wild-type (“reverse”) for each mutation. For the A99 protocol, ten repeated equilibrium simulations were used for charge-conserving mutations, and twenty for charge-changing mutations; from these, 400 frames were extracted for charge-conserving mutations, and 800 frames were extracted for charge-changing mutations. The non-interacting (“dummy”) atoms for morphing wild-type residues into mutants were introduced via pmx, using the mutant structure proposed by FoldX as a template.^48^ Positions of the dummy atoms were then minimized while freezing the rest of the system. These systems, now containing “hybrid” residues, were then simulated for 10 ps to equilibrate velocities.

Finally, amino acid side chains were alchemically morphed at constant speed during non-equilibrium simulations of 80 ps in length for A14 and 100 ps for A99. Work values associated with each non-equilibrium transition were extracted using thermodynamic integration (TI) and then used to estimate the free energy differences with the Bennett’s Acceptance Ratio (BAR).^53,129,130^ Point estimates of the free energy differences (ΔG_apo, forward_ and ΔG_holo, forward_) were calculated with BAR after pooling all available forward and reverse work values coming from the nonequilibrium trajectories. Uncertainties in ΔG_apo,forward_ and ΔG_holo, forward_ were estimated as standard errors (σΔG) by considering each equilibrium simulation and the resulting non-equilibrium trajectories as independent calculations. Uncertainties were then propagated to the final ΔΔG estimate to obtain the estimate of the standard error σΔΔG.

### Machine learning model deployment

The machine learning (ML) model was built in Python using the ExtraTreesRegressor class in the scikit-learn library, following the approach similar to Aldeghi et al., with variations in dataset splitting applied to feature selection procedures.^12^

#### Training dataset curation

The dataset described in Hauser et al. was used for training the model.^31^ This dataset contains 144 binding affinity changes (ΔΔG) for eight tyrosine kinase inhibitors (TKIs) due to point mutations in human Abl kinase. Six of these are structures resolved experimentally via X-ray crystallography (4WA9, 3UE4, 4XEY, 1OPJ, 3CS9, 3OXZ) and two were obtained via docking (referred to as DOK1 and DOK2).^91,103,131–133^ Models for mutant apo structures were generated using FoldX (v4). Structures of the mutant complexes were obtained by maintaining the ligand coordinates from the WT structures.

#### Features and feature selection

A total of 128 features spanning small-molecule and protein properties were considered as candidate model inputs: 18 ligand properties (e.g., molecular weight, calculated logP, number of rotatable bonds) were calculated with RDKit (v2018.09.1; https://www.rdkit.org), and 21 properties describing the mutation environment (e.g., distribution of ligand and protein atoms around the mutation site, number of polar/apolar/charged residues in the binding pocket) were calculated with Biopython (v1.73; www.biopython.org),^134^ 13 features describing the change in the amino acid chemical nature were calculated using precomputed properties for each amino acid (e.g., change in side-chain volume, hydropathy, number of hydrogen bond donors). Among these features, we also include the change in folding free energy upon mutation as predicted by FoldX v4. Six features describing protein-ligand interactions (hydrogen bonds, hydrophobic contacts, salt bridges, π-stacking, cation-π interactions, and halogen bonds) were calculated with the Protein-Ligand Interaction Profiler (PLIP).^135^ The Vina binding score, along with 59 Vina features were calculated with AutoDock Vina via scripts that are part of DeltaVina.^136^ The latter tool, in conjunction with the molecular surface calculation library MSMS, was also used to calculate 10 pharmacophore-based solvent-accessible surface area (SASA) features.

After candidate feature measurement was performed as described above, we performed a greedy algorithm using the ‘mlxtend’ library to select the most relevant features. During feature selection, we minimize the mean-squared error (MSE) of 10-fold cross-validation on the training set (a dataset of 144 ΔΔG values). Each of the ten folds were created such that each contained a unique set of mutations.

Feature selection was performed with a greedy algorithm using the ‘mlxtend’ library. We allowed the selection of any number of features up to a maximum of 40, which minimized the mean-squared error (MSE) of 10-fold cross-validation on the training set of 144 published ΔΔG values. The 10 folds were created such that each fold would contain a unique set of mutations. Using this protocol, we filtered our list of features down to the ten features that can best extrapolate to previously unseen point mutations. The ten features selected are: 1.

Change in hydrogen bond acceptor SASA between the WT and mutant complexes, 2. Change in halogen SASA, 3. Number of ligand-mutated residue atom pairs within 2 Å of each other (reporting on whether the mutation might introduce steric clashes), 4. Gain/loss of cation-pi interactions and salt bridges between the WT and mutant complexes, 5. Change in number of aromatic rings for the protein residue, 6. Change in the number of hydrogen bond acceptor atoms for the protein residue, 7. Maximal distance between ligand and WT residue atoms, 8, 9. The 25th and 50th percentiles of the values of the angles between each ligand atom and the beta and alpha carbons of the protein residue (a measure of the side chain orientation with respect to the ligand), 10. number of rotatable bonds in the ligand.

With these ten features, the model was then trained on the full set of 144 ΔΔG values from Hauser et al.^31^ This model was used to predict the ΔΔG values associated with the ABL1A kinase mutations described in this manuscript. The machine learning (ML) model was built in python using the ExtraTreesRegressor class in the scikit-learn library.^137^ This model uses ensembles of randomized decision trees in a similar fashion to random forest.^73^ The input files and the code (as Jupyter notebooks) used to train and test the ML models are provided online (https://osf.io/s6ktq/). All computations pertaining to the ML results were performed on a desktop machine equipped with an Intel Xeon processor of Broadwell architecture (E5-1630 v4). Splits were done by mutation. We split the Hauser dataset in 10 folds, such that each fold would have a different set of mutations, with the idea of encouraging the selection of feature that provide good extrapolation performance to new mutations (since we knew the ligands would be imatinib and dasatinib, but the mutations would be new).

#### Data analysis

The accuracy of the calculations was evaluated using three performance measures: the root-mean-square error (RMSE), the Pearson correlation (r), and the area under the precision-recall curve (AUPRC). The uncertainty in these measures was evaluated by bootstrap. Pairs of experimental and calculated ΔΔG values were resampled with replacement 105 times. For each bootstrap sample, RMSE, R, and AUPRC were calculated. From these 105 bootstrap measures, the 2.5 and 97.5 percentiles were taken as the lower and upper bounds of the 95% confidence interval. A bootstrap procedure was also used to obtain p-values for the differences between approaches. In this case, triplets of ΔΔG values were resampled with replacement together 105 times: ΔΔG values from experiment and from the two approaches to be compared. At each bootstrap iteration, the difference in the performance measure of interest (e.g. RMSE) between the two computational approaches to be compared was stored. At the end of the procedure, 105 bootstrap differences (e.g. ΔRMSE) would have been collected. The fraction of differences crossing zero was multiplied by two to provide a two-tailed p-value for the difference observed. Data analysis was performed in python using the numpy, scipy, pandas, scikit-learn, matplotlib, and seaborn libraries.^138–142^ All comparative analysis and summary statistics do not consider the mutations included in the dataset used to train the model, to reduce bias caused by data leakage.

#### Rosetta prediction of mutation impact using flex_ddg

Using the above protocol in the structure preparation and data curation of the PMX calculations, Rosetta binding free energy changes were calculated with Rosetta (v2017.52) using the flex_ddg protocol.^38^ These calculations were carried out on cluster nodes equipped with an Intel Xeon processor of S4 Broadwell architecture (E5-2630 v4), using one CPU core per ΔΔG calculation. Ligand parameters were obtained with the molfile_to_params.py script provided with Rosetta. The REF2015 and beta_nov2016 (referred to as βNOV16) scoring functions were used. The final ΔΔG estimates were the average values of the generalized additive model obtained from 35 iterations of the protocol. The command lines used for the Rosetta calculations and the input files can be found online (https://osf.io/s6ktq/).

#### NanoBRET affinity assay

Bioluminescence resonance energy transfer (BRET) is used as a proximity-based measure of drug binding to kinase targets in live HEK293T cells.^143^ BRET is observed between a NanoLuciferase (nLuc) tag on the full-length protein kinase and a tracer molecule (a BODIPY fluorophore attached to an ATP-competitive inhibitor scaffold). Upon binding of either imatinib or dasatinib to the kinase, the tracer is displaced to reduce BRET in a dose-dependent manner. The Abl NanoBRET affinity and residence time assay data used in this manuscript was collected as previously described by Lyczek et al.^24^ Briefly, full-length Abl was cloned in frame with an N-terminal NanoLuc fusion, mutagenized, and used to transiently transfect HEK293T cells at a density of 2x105 cells/mL for twenty hours. Transfected cells were incubated with BRET Kinase Tracer K4 (Promega) at the previously measured Tracer IC50 and serially diluted imatinib and dasatinib in OptiMEM media without phenol red. The system was allowed to equilibrate for two hours at 37°C and 5% CO2. The 3X Complete Substrate and Inhibitor solution was prepared by mixing NanoBRET Nano-Glo Substrate (Promega) and Extracellular NanoLuc Inhibitor (Promega) into OPTI-MEM. Tracer compound was added to cells using a liquid dispenser at the tracer IC50 concentration for each mutant,^24^ followed by addition of serially diluted inhibitor using an Echo 550 and incubated for 2 hours.

Afterwards, 3X Complete Substrate and Inhibitor were added to cells and luminescence was measured at 450 nm (donor emission). BRET was measured at multiple serially diluted concentrations of either imatinib or dasatinib and 650 nm (acceptor emission) in a PHERAstar plate reader (BMG Labtech). Background-corrected BRET ratios (610 nm/450 nm) were determined by subtracting the BRET ratios of samples from the BRET ratios in the absence of tracer and inhibitor. BRET ratios were plotted as a function of inhibitor concentration and graphed using GraphPad Prism (v9). IC50 determination was done via curve fitting to a four-parameter equation:

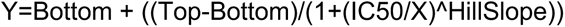

where the Hillslope describes the steepness of the curve (fixed to -1.0), and Top and Bottom describe the upper and lower plateaus in the units of the Y-axis, X is the ligand concentration, and Y is the BRET ratio. Apparent affinity values were each compound/mutant pair were calculated using the Cheng-Prusoff equation.^144^

### Retrospective analysis with Perses

#### Structure preparation and setup for alchemical transformations within Perses

Perses calculations were performed with structures of human wild type ABL1 in complex with imatinib and dasatinib prepared using functionality of OEChem and Spruce from the OpenEye Toolkits 2021.1.1 implemented in the open-source framework KinoML.^11^ PDB entries 2HYY (chain C) and 2GQG (chain A) were chosen as the ligands of interest were co-crystalized and due to the high quality score reported by KLIFS.^70,145–147^ Unresolved side chains of both structures were modeled with Spruce (OpenEye). The phosphorylated tyrosine of 2GQG at position 393 was altered to a standard tyrosine residue using Spruce. Missing residues of 2HYY, D276 and T389-D391 were built with Spruce. Finally, both structures were protonated at pH 7.4 using OEChem.This resulted in a charge of +1 for imatinib at the piperazine ring. Generated structures and scripts for structure preparation are available on github (https://github.com/openkinome/study-abl-resistance).

#### Perses hybrid topology setup

The hybrid topology, positions, and system for each transformation were generated using Perses 0.10.1 and OpenMM 8.0.0.^47,86,148^ The hybrid topology was generated using a single topology approach. The hybrid positions were assembled by copying the positions of all atoms in the WT (“old”) topology and then copying the positions of the atoms unique to the mutant (“new”) residue (i.e., unique new atoms). Unique new atom positions were generated using the Perses FFAllAngleGeometryEngine, which probabilistically proposes positions for one atom at a time based on valence energies alone. Further details on hybrid topology, positions, and system generation (including definitions of the valence, electrostatic, and steric energy functions) are available in the Perses “RESTCapableHybridTopologyFactory” class. For charge-changing mutations, counterions were added to neutralize the mutant system by selecting water molecules in the WT system that are initially at least 8 Å from the solute and alchemically transforming the WT water molecules into sodium or chloride ions in the mutant system. For example, if the mutation was ALA→ASP, a water molecule in the WT system was transformed into a sodium ion in the mutant system to keep the system endstate neutral. If the mutation was GLU→ALA, a water molecule in the WT system was transformed into a chloride ion in the mutant system. Additional details on the counterion implementation can be found in the Perses “_handle_charge_changes()” function found in “perses.app.relative_point_mutation_setup”. To prevent singularities when turning off the nonbonded interactions involving unique old or unique new atoms, a softcore approach was used that involves “lifting” unique old or unique new interaction distances: A padding distance (*w*(*λ*)) was added to the interaction distances involving unique old or unique new atoms so that the atoms could not be on top of each other.^149^ *w* lifting (the maximum value for *w*(*λ*)) was selected to be 0.3 nm. This lifting softcore approach was applied to both the electrostatic and steric interactions, so multi-stage alchemical protocols (e.g., where electrostatics must be turned off before sterics) were not necessary for scaling on or off the electrostatic and steric interactions. Instead, a linear protocol was used for interpolating the valence, nonbonded, and lifting terms. This softcore approach is similar to traditional softcore approaches with the critical difference being with our handling of the Lennard Jones potential. Our approach uses a lifting distance (*w*(*λ*)) that is independent of *σ* (the distance at which the Lennard Jones potential energy equals zero),^150,151^ whereas prior approaches define the lifting distance as a function of *σ*. In our approach, the lifting distance was defined to be independent of sigma for ease of implementation.

#### Replica exchange sampling using Perses (RepEx)

Alchemical replica exchange (AREX) simulations were performed using Perses 0.10.1 and OpenMMTools 0.21.5 (https://github.com/choderalab/openmmtools). The alchemical protocol was defined with evenly spaced *λ* values between 0 and 1. Before AREX was performed, the positions were minimized at each of the alchemical states using the OpenMM LocalEnergyMinimizer with an energy tolerance of 10 kJ/mol. Each AREX cycle consisted of running 250 steps (4 femtosecond timestep) with the OpenMM 8.0.0 LangevinMiddleIntegrator at a temperature of 300 K, a collision rate of 1 picosecond^-1^, and a constraint tolerance of 1e-6.^150–152^ All-to-all replica swaps were attempted every cycle.^51^ Replica mixing plots were created using OpenMMTools 0.21.5 (https://github.com/choderalab/openmmtools) to extract the mixing statistics from the AREX trajectories. Default settings were used unless otherwise noted. Full details on the AREX implementation are available online (https://github.com/choderalab/perses-b arnase-barstar-paper/blob/main/scripts/04_run_repex/run_repex.py). For each replica, 5000 cycles (i.e., 5 ns) were run, resulting in 30 ns of sampling per phase per mutation. Replicas mixed well for all mutations, indicating good phase space overlap. 10000 cycles were initially run per replica (10 ns/replica), resulting in 240 ns of sampling per phase per neutral mutation and 360 ns of sampling per phase per charge changing mutation. To improve the accuracy of our predicted free energy differences, the sampled alchemical states were bookended with “virtual endstates,” which were not sampled during free energy calculation, but for which reliable estimates of the physical endstates could be robustly produced during analysis.

#### System equilibration and alchemical sampling using Replica Exchange and Nonequilibrium cycling (NEQ)

All systems used the AMBER14SB force field for the protein and GAFF-2.11 for the small-molecule.^110,115^ Each system was then equilibrated for 3 nanoseconds (ns) where bonds to hydrogen were constrained with CCMA and Hydrogen Mass Repartitioning (HMR) with hydrogen masses set to 4 amu) was applied to allow for a 4 femtosecond (fs) timestep.^153^ A nonbonded cutoff of 1.1 nm was used for Lennard-Jones 12-6 interactions. Particle mesh Ewald (PME) was applied for treatment of long-range interactions with a direct-space cutoff of 1 nm, relative error tolerance of 0.0005, and automatic (default) selection of alpha and grid spacing.^125^ Alchemical Replica Exchange (RepEx) was performed on Perses using multiple swapping cycles to mix alchemical configurations.^47^ Each RepEx cycle consisted of running 250 steps (1 picosecond at a 4 femtosecond timestep) using the Leapfrog Langevin Integrator with a BAOAB splitting at a temperature of 300 K, a collision rate of 1 picosecond^-1^, and a constraint tolerance of 1e-6.^152,154^ All-to-all replica swaps were attempted every cycle. This set of cycles was repeated for every mutation in both complex (inhibitor-bound) and apo (inhibitor-absent) constructs of Abl kinase with both Dasatinib and Imatinib complex structures.

Replica mixing plots were created using OpenMMTools 0.21.5 (https://github.com/choderalab/openmmtools) to extract the mixing statistics from the AREX trajectories. Default settings were used unless otherwise noted. Full details on AREX implementation are available on github (https://github.com/choderalab/perses). Non-equilibrium cycles (NEQs) are independently collected on Folding@home where each individual cycle serves as an independent statistical replicate in free energy estimation.^39,95,155^ A single cycle consisted of 4 stages each of which lasted 1.5ns. An initial equilibration at λ = 0 was first run, followed by a forward non-equilibrium process which drives λ from 0 to 1 over 100 equally spaced windows, followed by another equilibrium simulation at λ = 1. Lastly, a reverse non-equilibrium process driving λ from 1 to 0 was run over 100 equally spaced windows across another 1.5ns. All cycles were run using a Leapfrog Langevin Integrator with BAOAB splitting at a collision rate of 1 picosecond^-1^ and a constraint tolerance of 1e-8, leading to a total trajectory time of 6ns per cycle. 100 replicate cycles were collected per mutation in both complex (inhibitor-bound) and apo (inhibitor-absent) on Folding@home to obtain forward and reverse work distributions. For both RepEx and NEQ sampling using Perses, free energy differences were calculated using the Multistate Bennett Acceptance Ratio.^156^

#### Equilibrium simulations of T315 and T315A to observe water coordination

Starting from the same structure as was used for Perses NEQ trajectories and bound to imatinib, as described above, we define a hydrated starting structure of Abl kinase as hydrated if the closest water molecule is within 0.4 nm of residue 315. We then generate the dehydrated starting structure by identifying the single water molecule within 0.4 nm of residue 315 and delete it. Both of these structures, in WT and mutant forms, are then used as the starting structures for equilibrum MD simulations using Gromacs 2016. For both hydrated and dehydrated starting structures, energy minimization was carried out using a steepest descent algorithm for 1000 steps. The systems were subsequently simulated for 100 ps in the isothermal-isobaric ensemble (NPT) with harmonic position restraints applied to all solute heavy atoms with a force constant of 1000 kJ mol^-1^ nm^-2^. Equations of motion were integrated with a leap-frog integrator and a time-step of 2 femtoseconds (fs). The temperature was coupled with the stochastic v-rescale thermostat at the target temperature of 300 K.^120^ The pressure was controlled with the Berendsen weak coupling algorithm at a target pressure of 1 bar.^121^ The particle mesh Ewald (PME) algorithm was used for electrostatic interactions with a real space cut-off of 10 Å when using Amber force fields, a spline order of 4, a relative tolerance of 10^−5^, and a Fourier spacing of 1.2 Å.^122^ Verlet cut-off schemes with the potential-shift modifier was used with a Lennard-Jones interaction cut-off of 10 Å, and a buffer tolerance of 0.005 kJ mol^−1^ ps^−1^.^123^ All bonds were constrained with the P-LINCS algorithm.^124^ For equilibration, unrestrained MD simulations were then performed for 1ns in the NPT ensemble with the Parrinello-Rahman barostat at 1 bar with a time constant of 2 ps.^125^ Production simulations were then performed for 100 nanoseconds for both hydrated and dehydrated structures. The above was then repeated for both wild type (with T315) and mutant T315A, where the T315A mutant structure was extracted from Perses NEQ trajectories generated above at the mutant endstate (where λ = 1).

## RESULTS AND DISCUSSION

### NanoBRET is consistent with previous experimental measurements of mutational impact to identify inhibitor-resistant and -sensitizing mutations

In an effort to understand the clinical relevance of mutations within the ABL1A kinase domain, we gathered mutations observed in clinical settings using known databases and quantified mutation prevalence across clinical samples (Fig. 1).^5^ While there are multiple possible mutations per position, the total number of clinical samples with a particular mutation can be represented using a lollipop plot (Fig. 1B). Mapping the distribution and frequency of clinical mutations to the structure of ABL1A kinase domain (Fig. 1A) also allows us to identify structural hotspots with high mutation propensity, or pinpoint hotspots where mutations are particularly prevalent in ABL1A. We note that mutations are distributed throughout the secondary structure and sequence of ABL1A, and while certain positions have many mutations, mutations do not appear to concentrate to any particular hotspot or secondary structural kinase element.

We obtain NanoBRET measurements that quantify the impact of 90 highly prevalent mutations from this clinical dataset (Fig. 2, SI data). NanoBRET allows for precise quantification of the changes in drug binding affinity caused by these mutations.^24,68,69^ These NanoBRET measurements measure the degree of inhibitor-target engagement using bioluminescent probes that luminesce when bound to one another.^24,68,69^ These engagement measurements can be converted into an IC50 by scanning engagement across a variety of titratable concentrations.^24^ Our structural map demonstrates the broad distribution of mutated residues in relation to the drug binding site (Fig. 1A)

We can also assess the fold-change in IC50 for any mutant’s imatinib- or dasatinib-affinity relative to wild type (WT). Mapping the maximal fold-change onto the three-dimensional structure of ABL1A kinase (Fig. 2A) allows us to visualize the range of impact mutations have on ligand binding. Mapping the maximum fold impact of a mutation at a specific position from NanoBRET onto the structure of Abl kinase highlights the broad distribution of mutated residues relative to the inhibitor binding site (Fig. 2A). Notably, the color-coded mutations illustrate a spectrum of impacts, with some mutations causing increased inhibitor resistance, while others surprisingly lead to increased inhibitor sensitivity. The latter suggests that certain mutations, rather than conferring resistance, render the kinase more amenable to inhibition by existing drugs, thereby identifying potentially promising targets for therapeutic intervention. Mutations found to induce inhibitor sensitization may present promising targets for existing drugs, while others require additional treatment development.

We compare these NanoBRET measurements with previously published work on ABL1 mutations and their impact on imatinib and dasatinib binding affinity (Fig. 2B).^31,58,60,61^ By comparing the IC50 of wild-type and mutant ABL1A binding to a known inhibitor, imatinib or dasatinib, we estimate the change in free energy of binding (ΔΔG); the NanoBRET protocol using tracer concentrations at the K_M_ ensures that significant changes in IC50 will be largely driven by changes in inhibitor binding affinity. The ΔΔG values from previous works can be correlated directly against these ΔΔG measurements (Fig. 2B). The comparison of ΔΔG values obtained from NanoBRET with those from prior experimental data exhibits a noteworthy consistency for both Imatinib and Dasatinib. This consistency is quantitatively supported by low RMSE values and decent correlation coefficients (RMSE=0.61 kcal/mol for imatinib, and 0.59 kcal/mol for dasatinib), indicating the accuracy of our NanoBRET assay. It is worth noting that our NanoBRET measurements are derived from single-shot experiments, ensuring a high-throughput collection protocol. As such, we estimate error associated with each NanoBRET from the goodness-of-fit for each individual mutation (Fig. 2B, SI data). This provides a nuanced estimate of the experimental uncertainty despite the lack of statistical replicates. In conclusion, low RMSE and correlation scores not only reinforce the consistency of the NanoBRET measurements with previously published results but also underscore NanoBRET’s utility as a tool for the high-throughput screening of kinase mutations.

### Free energy calculations prospectively predict the change in affinity with RMSE of 1 kcal/mol and provide a robust classification metric for predicting a resistant or sensitizing mutation

While NanoBRET is a robust high throughput tool to assess mutations and their impact on inhibitor binding, it would be particularly useful, and cost-effective, for computational approaches to *prospectively* parse mutations as resistant, sensitizing, or neutral relative to any particular inhibitor. By forecasting the impact of genetic alterations on drug efficacy, computational predictions promise a preemptive tailoring of therapeutic strategies.

Prospective testing is crucial to verify the applicability of computational models. Retrospective analyses may inadvertently incorporate biases from known outcomes and contain outdated information and biases influence benchmarking comparisons. A prospective approach evaluates the predictive power of models in novel, untested scenarios, where tools are employed in standard default fashions without any biased use of the tool to a known outcome. This forward-looking methodology is essential for validating the robustness of predictive algorithms and ensuring their utility in clinical settings.

To effectively classify mutations as inhibitor-resistant or -sensitizing, a variety of computational and machine learning approaches were prospectively tested against NanoBRET. These methods include free energy calculations using two different sampling strategies: Hamiltonian Replica Exchange (RepEx or HREX),^47,49–52^ and Non-Equilibrium perturbations (NEQ),^12,31,36,47^ Rosetta flex_ddg,^12,38^ and a machine learning model utilizing Random Forests (referred to as “Random Forest”, “ML”, “RF”, or “AI/ML” here).^10,12^ Each method offers a unique computational strategy to predict the impact of mutations on protein stability and drug binding (Fig. 3).

**Figure 3.**
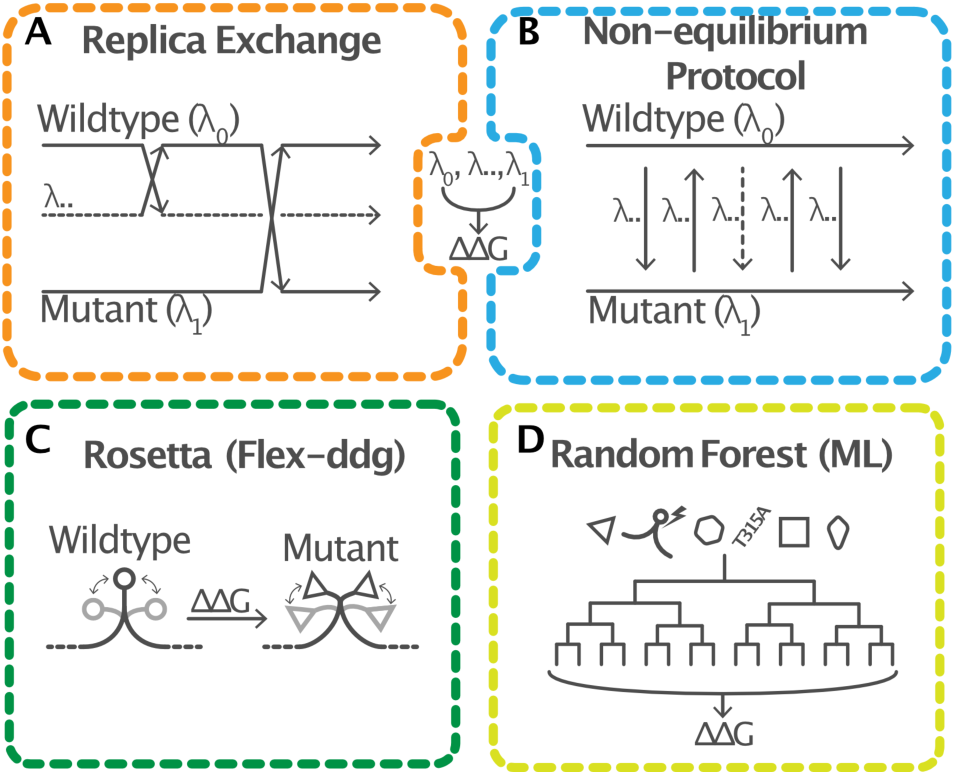
Overview of computational methods used to prospectively assess estimation of ΔΔG upon mutation. Physics-based alchemical methods (top) estimate the impact of mutation on drug binding (ΔΔG) by generating an alchemical transformation between wild-type (WT) and mutant constructs using a lambda parameter sampled using **A.** Replica Exchange methods and **B.** Non-equilibrium protocols. **C.** The Rosetta flex–ddg protocol considers rotameric positions of both WT and mutant amino acid side chains, using the Rosetta scoring function to compute the energetic impact of both constructs. **D.** Random forest models generated using an ensemble of decision trees are trained on prior assay measurements and structural data to generate ΔΔG value predictions.^31^

AI/ML approaches that directly assess the impact of mutations on drug binding have the potential to reduce the computational cost of predicting drug resistant mutations. In fact, many powerful models have been published that accurately predict the impact of mutations on protein stability.^74–77^ While stability is correlated with drug binding, it would be particularly useful if models provided direct readouts of the impact upon drug binding. In this work we will focus on prior published models that directly predict ΔΔG upon mutation of drug binding; Specifically, we will focus on a prior published Random Forest model to assess Abl kinase mutations.^10,12^ The model was trained on a curated dataset of 144 ΔΔG values for eight tyrosine kinase inhibitors, incorporating a diverse set of features that describe ligand properties, mutation environments, amino acid changes, protein-ligand interactions, docking scores, and solvent accessibility.^31,58,60,61^ Feature selection was performed using a greedy algorithm to identify the most predictive features.

Comparing each method against known mutations that have been previously studied highlights the capacity of each method to reasonably predict the impact of mutation on imatinib or dasatinib affinity. To test the prospective accuracy of these methods, we compared their predictions against experimental NanoBRET measurements for mutations in a manner similar to previous studies (Fig. 4).^31,58,60,61^ We observe that many prospective measurements capture a similar range of ΔΔG predictions from -6 to +6 kcal/mol, with the vast majority of ΔΔG predictions ranging between -2 to +2 kcal/mol (Fig. S10). Our comparative analysis demonstrates that while each approach has varying degrees of success, they collectively exhibit a reasonable predictive capacity for the same set of mutations.

**Figure 4.**
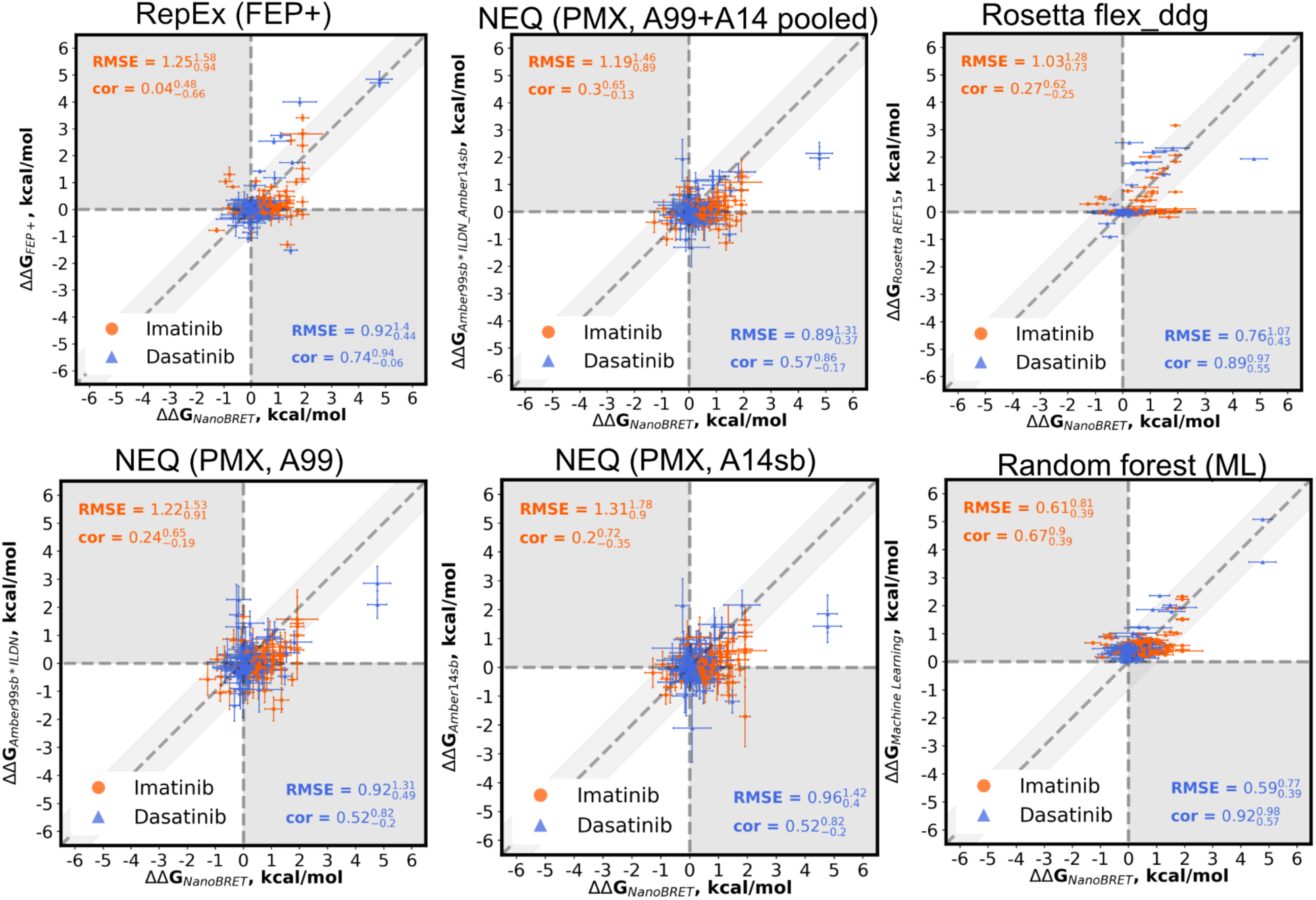
Prospective computational methods have comparable summary statistics and performance in predicting the impact of mutation on binding. Scatterplots showing the prospective ability of different computational methods and their ability to predict the ΔΔG of either imatinib (orange) or dasatinib (blue) binding. Root Mean Square Error (RMSE) and Pearson correlation (labeled “cor”) are provided in the top left (imatinib) and bottom right corners (dasatinib). These comparisons are all done relative to the same ΔΔG measurements collected using NanoBRET. Prospective methods shown are Replica Exchange using FEP+ (top left), Nonequilibrium switching using PMX using Amber99 force field (bottom left), the Amber14sb force field (bottom middle), and the resultant prediction taken from pooling the work values from both force fields (top middle). Rosetta’s flex_ddg (top right) and a random forest model trained on prior data (bottom right) are also shown.

Interestingly, all approaches appear to consistently do better at predicting the impact of mutations on dasatinib binding than imatinib binding, highlighted by the lower RMSE for dasatinib vs. the RMSE of imatinib across all the approaches (Fig. 4). This is further supported by the degree of correlation between calculated changes in affinity and NanoBRET measurements in the datasets. Interestingly, physics-based methods consistently predict the impact of mutations on dasatinib more accurately than imatinib. This difference in predictive capability for physics-based simulations may be related to the degree of conformational sampling required to accurately sample imatinib binding vs. dasatinib binding. Physics-based simulations are unique in their dependence on conformational sampling, something that has no impact on data-driven methods such as the Random Forest model. Previous work has shown that to characterize the complete ABL1 imatinib binding, a conformational transition in ABL1A must occur.^78–80^ However, efforts to characterize the full ensemble of structural transitions on binding have been limited by sampling, and there may be additional ABL1-Imatinib configurations.^26,81,82^ No such conformational transition or degree of sampling is needed to sample dasatinib binding.^78^ As such, there may be currently unseen states of the ABL1-Imatinib bound complex that must be considered in addition to the singular crystal that was used as a starting configuration here. The differences between dasatinib and imatinib are consistent across the different MD force fields as well as Rosetta. Thus, it may be that each ligand requires different amounts of conformational sampling during these calculations to achieve an average deviation of 1 kcal/mol from the NanoBRET measurements.

Marking the 1 kcal/mol boundary as an acceptable margin of error and degree of significance for perturbative mutations based on prior established work,^46,83–85^ we can categorize mutations as either resistant, sensitizing, or neither (Fig. S2). Previous results show that most reproducible estimations occur with an RMSE of up to 1 kcal/mol.^12^ To then capture the most reproducible predictions, we consider a mutation to be correctly assigned as resistant if both the NanoBRET and computational approach predicted an increase in ΔΔG by ΔΔG > +1 kcal/mol for a single inhibitor. Conversely, a mutation is considered to be correctly assigned as sensitizing if both NanoBRET and computational predictions identify drug-binding affinity is decreased by ΔΔG > -1 kcal/mol. We can visualize these classifications as “quadrants” in our correlation analysis, generating truth tables (confusion matrices) for each computational method (Fig. S1).^31^ This matrix allows us to evaluate the performance of each computational approach in correctly classifying mutations and compare that performance statistically against a baseline classifier.

From these truth tables, we show that computational approaches classify mutations as resistant or sensitizing better than a baseline classifier would, shown using a precision recall curve (Fig. 5, Table S1). A precision-recall curve graphically represents model performance in contexts where the classifications have different populations and frequencies. This allows us to evaluate the sensitivity and specificity of our measurement and model. Precision, known as positive predictive value, measures the ratio of the predictive model’s true positives to all the positive predictions. Recall, also defined as sensitivity, measures the model’s ability to identify all true positives from the data. This allows us to identify the best threshold for the predictive model, and helps us establish whether our classifier is performing better than random behavior. A higher area under the curve (AUC) indicates a better model performance. We compare the performance of each computational approach as a classifier to a baseline classifier (Fig. 5, Table S1), where the baseline classifier reports the performance if everything was labeled as statistically positive, or in our case, a resistant or sensitizing mutation. For the 1 kcal/mol cutoff, all approaches perform better than a baseline classifier, having a positive distance above the baseline as shown by the positive distance above the baseline (Table S1).

**Figure 5.**
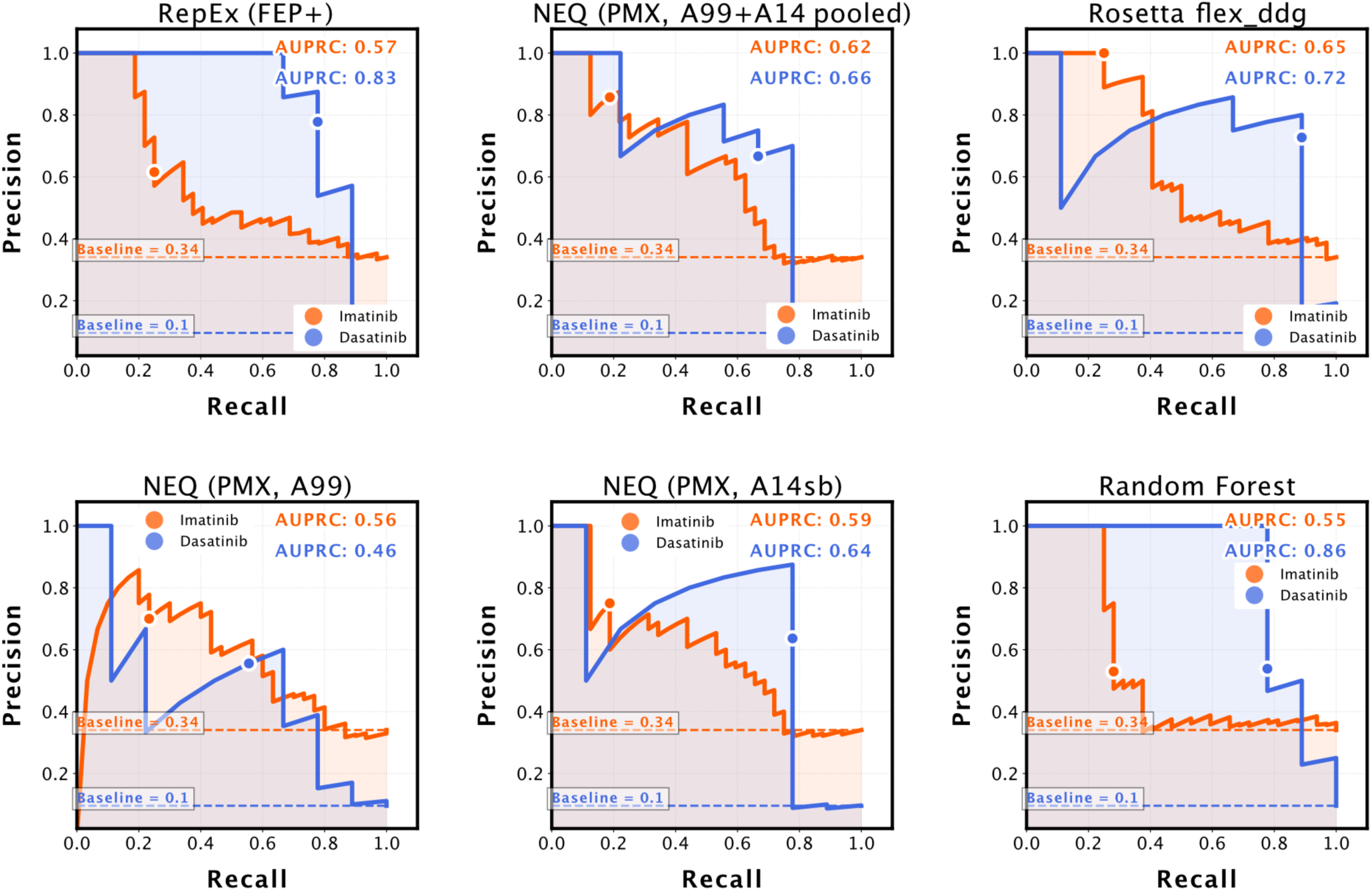
Precision-Recall Curves for each method to demonstrate their ability as classifiers is better than random, and that they are all similar in performance. Precision is defined as the fraction of mutations that are classified as resistant that are actually resistant, while recall is the fraction of mutations that are classified as resistant. The area under the precision recall curve (AUPRC) is computed for both imatinib (orange) and dasatinib (blue). The singular point on the curves corresponds to the precision (x) and recall (y) value for mutations with |ΔΔG| > 1 kcal/mol, and are computed for imatinib (orange) and dasatinib (blue), and compared against the performance of a random classifier (dashed line).

The distance from the baseline at 1 kcal/mol acts as a useful measure to compare both the performance of each approach against one another as well as between the two different drugs. At the 1 kcal/mol margin, it is worth noting that the Random Forest model is closest in performance to the baseline classifier (Table S1), indicating that it performed relatively worse than the other methods as a classification tool compared to a statistical baseline. The larger number of false positives (mutations that computationally are not predicted to reduce the affinity but experimentally do) indicates that the Random Forest model was over-assigning resistant/sensitizing classifications. Consistent with RMSE metrics, we find that imatinib is much harder to classify relative to baseline than dasatinib for many of the tools, with each approach having a much greater distance to baseline when classifying dasatinib than imatinib. Our Random Forest model is particularly bad at classifying imatinib mutations relative to dasatinib, with its overall performance being worse than the physics based alchemical method. While Random Forest and Rosetta-based predictions demonstrate similar performance to alchemical methods, with only a slightly worse performance as indicated by the increased number of false positives, it is worth noting that these are cheaper to run computationally by orders of magnitude.^47,86^ Historically, Alchemical methods are computationally expensive models, with sufficient sampling requiring orders of magnitude more computational time than Rosetta. Meanwhile Rosetta and Random Forest models can scan across multiple predictions in much smaller time. Random Forest models, in particular, are well suited for rapid prediction of mutational impact and can be deployed in a few lines of python code on commodity hardware once training is complete. Given their similar performance to physics-based methods, data driven methods like Random Forest models may be well suited to rapidly pare down candidate mutations proximal to the inhibitor for extended study.

Taken together, these results highlight the capacity of the employed computational methods to act prospectively as classifiers within an average accuracy of 0.81 for resistant or sensitizing mutations (Table S1). Modern computational approaches act as useful classifiers for sorting clinically observed mutations into useful, potentially actionable categories. These approaches appeared very similar to one another using RMSE and correlative measures, with no clear indicator of relative performance. We can compare methods across multiple inhibitors, identifying performance improvements as well as points of failure. This comparison is achieved through the use of precision-recall curves, computing the area underneath precision recall curves (AUPRC), and comparing each method to a baseline classifier. Critically, our assessments using Precision-Recall curves highlight the importance of analyzing prospective predictions beyond computing summary statistics and correlative measurements, as these ensemble findings may hide key differences.

### Alchemical methods can detect impact of distal mutations upon ligand binding

While each of these methods may be similarly capable of predicting the impact of mutations on binding, summary statistics and analysis may mask issues when considering prospective benchmarking predictions. In turn, it is important to consider each mutation’s prediction individually to observe similarities or performance improvement. For example, the similarity of results shown from the classifier model may mask a degree of per-residue and per-mutation variance that results from a larger systematic issue.

By looking at the maximum possible improvement possible by simulation per mutation, we find that, while predictions are within ∼1 kcal/mol experiment, free energy calculations generally offer more improved predictions at any position in the sequence. By subtracting the least accurate non-free energy calculation prediction (Random Forest/ML and Rosetta) from the most accurate alchemical prediction, we compute the maximum amount that physics-based simulations are capable of making better predictions (Fig. S3). A more negative improvement score thus conveys that simulations were much closer to the answer than non-free-energy methods were, while a positive score indicates the reverse. We find that, for any single mutation, free energy calculations provide a small but consistent margin of improvement, at most 0.5 kcal/mol, relative to alternative methods such as Rosetta or ML-based methods (Fig. S3).

Potential improvements offered by simulation-based methods such as alchemical physics-based simulations is further emphasized when considering allosteric mutations (i.e. distant residues relative to the ligand binding). Identifying distal mutations that perturb binding free energy remains a challenge for modern computational methods but has great potential to help prospectively identify allosteric mutations and cryptic pockets.^23,40,41,87–90^ However identifying significant distal mutations, and prospectively analyzing them, remains a challenge due to their relative infrequency in clinical datasets that identify significant mutations with known impact.

We consider how distal a mutation is relative to the ligand binding site by computing the distance from the center of mass of a residue to the center of mass of the inhibitor. We first note that experimental ΔΔG from NanoBRET experiments show many distal mutations impact inhibitor binding, with residues up to 30 Å away having an impact on ΔΔG of imatinib binding (Fig. 6A). However, we note that many residues have large error bars due to the singlicate measurements. A significant alteration to ΔΔG would be much clearer if a mutation with ΔΔG is non-zero while its error bars are much smaller (Fig. S4–S6). Most residues with significant impact on inhibitor binding are in closer proximity to the binding site, with only a few distal mutations having significant impact (Fig. 6A, Fig. S5, S6).

**Figure 6.**
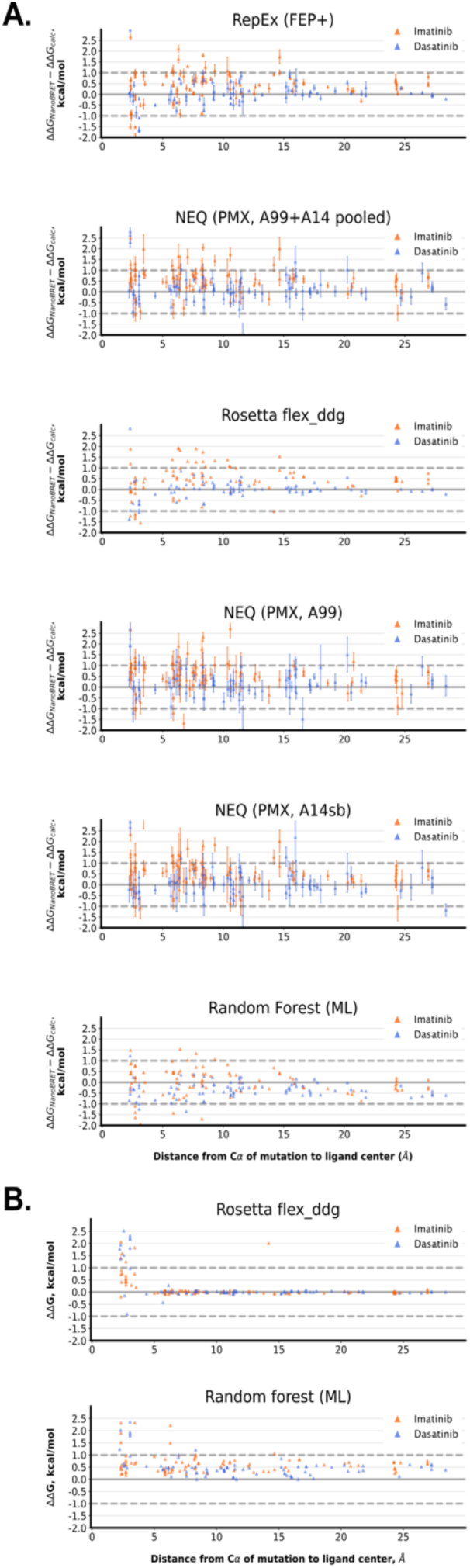
Physics-based methods are capable of estimating ΔΔG for distal mutations, while default parameters for Rosetta and ML-based methods collapse at a distance. **A.** The deviation from predicted ΔΔG to experiment is plotted for imatinib (orange) and dasatinib (blue) as a function of the distance from the residue’s Cα carbon to the center of mass of the ligand in the crystal structure. Values for shown (top to bottom) for Replica Exchange using FEP+, PMX predictions taken from pooling the work values from both Amber99 and Amber14sb, Rosetta’s flex_ddg protocol, Nonequilibrium switching using PMX using Amber99 force field, followed by Nonequilibrium switching using the Amber14sb force field, and a random forest model trained on prior data **B.** Predicted ΔΔG values for shown (top to bottom) for Rosetta’s flex_ddg protocol and the random forest model trained on prior data, indicating the model predicts within a small range of values for distal mutations.

By projecting the predicted ΔΔG or the deviation of predicted ΔΔG from experiment (Fig. 6B, Fig. S6), we note that both Rosetta and the Random forest (ML) methods appear to have systematic biases in their predictions (Fig. 6B); This is especially obvious for Rosetta and ML methods when looking directly at the ΔΔG predicted values as a function of distance. Both approaches appear to flatten out and center around a single value, with Rosetta predictions returning ΔΔG=0 kcal/mol beyond a certain distance from the active site. Similarly, the random forest model makes predictions for residues within the binding pocket of imatinib beyond 1 kcal/mol, collapsing to predictions around a ΔΔG∼0.5 kcal/mol upon moving further away from the drug binding pocket. This systematic error is likely the result of in-built distance cutoffs in the computational approaches. By enforcing a cutoff to make mutation ΔΔG predictions computationally tractable, a mutation’s local environment is mainly considered. As a result, while a mutation may have some impact on its local and global environment, constraining only to local effects will ignore any impact a distal mutation may have on inhibitor binding affinity.

Conversely, we note that free energy calculation predictions appear to deviate from experiment more randomly, but remain within 1 kcal/mol even at a distance (Fig. 6A, Fig. S4). Consistent with experimental results, residues predicted to have the strongest impact on ΔΔG are generally closer to the active site.

Importantly, even at a distance, the vast majority of computed deviation from experimental values appears to be below 1 kcal/mol (Fig. 6A). However, due to the error within the NanoBRET experiments, it remains unclear if this low margin of error is computationally driven, or if these mutations themselves do not impart a very large ΔΔG upon inhibitor binding, reducing the possible margin of error. Regardless, alchemical methods appear capable of predicting and considering the impact of distal mutations.

Structural considerations on a per-mutation basis allow us to more broadly assess the prospective ability of computational methods to predict the impact of mutation on an inhibitor’s binding affinity. The error bars within our experimental results also further highlight the importance of considering experimental error as well as computational error when considering predictions.

### Retrospective analyses of clinically significant mutations emphasize the importance of starting configurations, protonation states, and sampling

While each method appears to predict the impact of each mutation to a consistent rate, based on summary statistics and AUPRC findings, it is interesting to also note the consistent points of failure. Certain clinically notable mutations such as T315A, a known clinical mutation that causes imatinib resistance,^16,91,92^ are inaccurately predicted in a consistently poor manner. In fact, PMX based NEQ protocols are unable to predict even the correct sign of T315A (Table S2). The consistent failing to accurately predict the ΔΔG of certain mutations suggests a systematic error across all methods that results in this inaccurate prediction.

There are many possible sources of error that could systematically bias ΔΔG estimations. Indeed, considerable effort has gone into characterizing the source of errors that occur in ΔΔG predictions from replica exchange alchemical transformations.^47^ These previous efforts identified many potential system specific structural features that might be slow to converge in a replica exchange sampling schema. Consistently, there may be structural features that are slow to converge that are the basis of some errors in the alchemical transformations. However, other ΔΔG estimations presented here do not necessarily use physics-based simulations, instead utilize other information, sampling schemes, or alternative datasets to generate their predictions. As a result, there are only a few common systematic sources of biases due to the variable parameters and defaults used in each estimation. The consistency in deviation from NanoBRET across multiple approaches may also hint at discrepancies between experimental measurements of target engagement and the direct binding affinity estimated via computation (Table S2).

To retrospectively analyze these predictions in an open-source, interpretable manner at scale, we employ the open source free energy tool, Perses.^47^ The Perses toolkit, built on the OpenMM engine, is an open-source scalable codebase that allows us to consistently assess each mutation. The flexible architecture of the toolkit also enables us to try multiple simulation parameters during retrospective analysis. In doing so, Perses enables us to flexibly assess different simulation parameters and their impact on ΔΔG prediction accuracy.

Open source tools, Perses and PMX, allow us to assess multiple sampling strategies, considering both replica exchange and nonequilibrium methods using a single unified implementation (Fig. S7, S8).^48,93,94^ To quickly generate ΔΔG predictions using consumer-grade GPU architectures, we leverage the Folding@home distributed computing platform to run our ΔΔG predictions. By running on consumer-grade GPUs, our simulations can run on any type of GPU system, maximizing reproducibility (Data availability statement). For example, we can rapidly generate NEQ estimations using Perses in a massively parallel manner by deploying alchemical simulations of mutations on Folding@home.^39,95^ Folding@home is a distributed computing platform where molecular simulations are run in discrete work units. The asynchronous nature of each work unit makes Folding@home particularly suited to evaluate many mutations in parallel.^37,96^ Indeed, prior computational work has leveraged Folding@home’s massively parallel nature to evaluate mutations and potential inhibitors in an embarrassingly parallelizable manner using NEQ sampling; each work unit can run an individual replicate of a singular forward and reverse transformation.^37,96,97^ These individual “cycles” of NEQ replicates can then be aggregated to generate ΔΔG estimations (Fig. S6, S7). To obtain detailed assessments of the source of biases within our estimations, we focus on emblematic examples of clinically relevant mutations that were inaccurately predicted. Specifically, we consider two mutations: the perturbative “gatekeeper mutation” T315A mutation with known clinical significance to give rise to imatinib resistance,^16,91,92^ and the known stabilizing mutation L298F that improves imatinib binding affinity and increases sensitivity (Fig. 7, Table S2–S4).^24^ Identifying sources of errors spanning both these mutations, one perturbative and one stabilizing, can suggest generalized sources of errors that span different mechanistic impacts.

**Figure 7.**
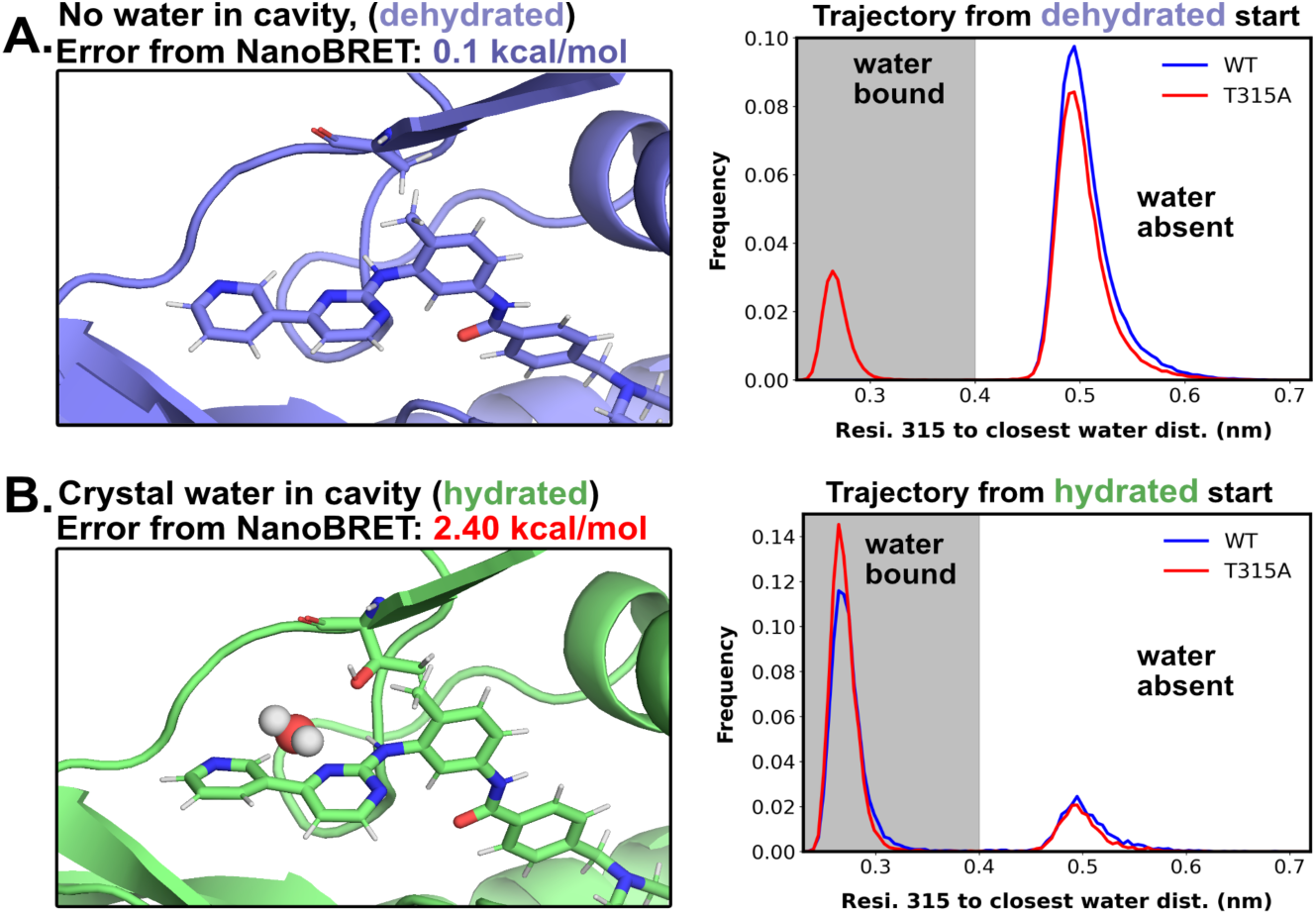
Starting configuration details such as water placement, play a major role in ΔΔG accuracy. **A.** Structure of a dehydrated T315 residue (left, blue) that yields a low error in ΔΔG prediction between physics-based simulations from Perses and PMX (below image). Equilibrium MD trajectories (top right) started from this dehydrated structure for both WT (red) and the T315A mutant (blue) show that the closest water molecule rarely comes within 4 Å of the residue (grey region), largely keeping the residue away from water (white region). **B.** Conversely, structures started from a hydrated structure (bottom left, green), where the closest water molecule is within 4 Å of residue 315, yields highly inaccurate ΔΔG predictions relative to NanoBRET (below image). Consistently, equilibrium MD trajectories (bottom right) started from the hydrated structure for both WT (red) and T315A (blue) show that the water largely remains bound (grey region) to the molecule for the same number of steps that the alchemical free energy predictions are run for, with little water dissociation occurring (white region).

Perses retrospective ΔΔG predictions are consistent with prospective predictive findings, showing that it is a useful platform for benchmarking and testing our results (Fig. S7, S8). Perses predictions appear to be consistent in summary statistics with findings from both RepEx and NEQ (Fig. S7). Perses predictions appear to behave similarly to other approaches when considered as a classifier (Fig. S7), returning equivalent or better precision-recall statistics and F1 scores (Fig. S7), and are similar to FEP+ and PMX (A99 + A14 pooled) predictions. Retrospective calculations using Perses also appear to capture the impact of distal mutations by FECs with a similar degree of consistency to other free energy methods (Fig. S8). Deviation of predictions by Perses from experiments across distances also appears similar (Fig. S8); This consistency with other free energy predictions demonstrates that Perses provides an appropriate platform to conduct detailed testing of high error estimations. Importantly, improvements made in Perses ΔΔG predictions would generalize and apply to other platforms and tools as well.

In the case of T315A, we find that the systematic failure in ΔΔG prediction is corrected by alteration of the starting configuration used to run these free energy calculations (Fig. 7, Table S2–S3). In our estimation attempts with T315A in the presence of imatinib, we find that ΔΔG estimations are highly inaccurate; Both physics-based sampling methods, RepEx and NEQ, fail to predict the correct sign of ΔΔG upon the T315A transformation (Fig. 7, Table S2–S3). Given this broad inability across multiple methods to accurately predict the correct sign of ΔΔG, the source of consistent error is likely related to some shared input or parameter that drives this error. In turn, we investigated the starting structure provided for Abl-imatinib for running ΔΔG predictions, since that is a common starting point for each of these protocols.

We observe a water molecule adjacent to T315 that we conjecture is having nonbonded interaction with the polar side chain of threonine (Fig. 7B). However, upon mutation to alanine, one would expect that the water would exit that interacting region due to the lack of interaction. We investigate the exchange of water within this pocket by measuring the distance between residue 315 and the closest water molecule at each frame (Fig. 7).

To better probe the role this water plays in the accuracy of our ΔΔG predictions, we must first define a structure where a water is present (Residue 315 is “hydrated”) and where a water is absent (residue 315 is “dehydrated”). Once we have appropriately defined a “hydrated” and “dehydrated” condition for residue 315, we can then extract representative structures with which to run additional simulations. From these simulations, we can compare how ΔΔG changes between the hydrated and dehydrated structures, informing the importance of this water around residue 315.

Based on these structures, we define a hydrated starting structure of Abl kinase as hydrated if the closest water molecule is within 0.4 nm of residue 315. Investigating the hydration status of residue 315 in Perses-NEQ trajectories reveals that the closest water molecule never dissociates about 5 Å away from the binding site (Fig. 7). To test whether this hydrated state is consistently present at equilibrium, we ran equilibrium MD simulation from the hydrated starting structure with both T315 and A315 (Fig. S9). We observe that the closest water molecule rarely dissociates away from residue 315 (Fig. S9). While a few transitions are observed where the water molecule dissociates from residue 315 (Fig. S9), we hypothesize that this lack of water dissociation upon the T315A mutation may be a source of error.

Consistently, we find that ΔΔG estimations that start from dehydrated structures, where the closest water molecule to residue 315 is deleted, improves our predictions (Fig. 7A). Using these dehydrated structures as starting configurations in both Perses and PMX structures, the sign of our predictions aligns with experimental results and reduces the deviation from experiment to 0.10 kcal/mol (Fig. 7A). Consistently, trajectories starting from a dehydrated starting structure only rarely observe a water molecule come within 4 Å of the mutated T315A construct, but a water molecule still does associate with the wild type T315A simulations (Fig. S9).

Interestingly, both simulations starting from dehydrated structures have a degree of mixing; water in the dehydrated trajectories are both interacting with T315 and dissociating (Fig. 7, Fig. S9). Overall, these findings indicate to us that in the absence of an appropriate starting structure, sufficient sampling of water configurations would enable accurate predictions. As a result, we recommend ensuring sufficient sampling using established metrics.^98,99^ As well as using a long enough switching time both to ensure appropriate structural relaxation to enable equilibration of solvent molecules.

For both T315A and L298F, we find that considering alternative protonation states improves ΔΔG estimations (Table S2–S4). Due to resource limitations and default parameters, prospective methods only consider a singular protonation state, imatinib with 0 net charge, which we call imatinib+0. However, imatinib can also exist at physiological pH conditions with a +1 charge, a species we call imatinib+1. While dasatinib can be protonated, at physiological pH there is likely to be only one charge species present in solution.^78,79,100^ As such, dasatinib does not have multiple likely titratable states, and so at physiological pH only one species needs to be considered.

For T315A and L298F, prospective methods struggle to obtain consistent ΔΔG predictions for imatinib+0 (Table S2–S4). Some methods such as FEP+ estimate the correct sign of ΔΔG upon the L298F mutation relative to the experimental NanoBRET measurement (Table S4). However, prospective evaluation of T315A is much more difficult for all three prospective methods, FEP+, PMX with A99 force field, and PMX with the A14 force field (Table S2, S4). Prospective ΔΔG measurements are unable to get the correct ΔΔG sign when predicting the impact of the T315A mutation on imatinib+0 binding (Table S2, S4).

We find that considering both protonation states of imatinib improves prediction capabilities. Running Perses RepEx methods or NEQ on Folding@home, we estimate ΔΔG for both T315A and L298F mutations for both imatinib protonation states. Using previously established approaches to consider the bulk ΔΔG given multiple protonation states,^47^ we compute a ΔΔG estimation in the presence of “bulk” imatinib, considering both protonation states by their propensity to exist at a given pH. Considering both protonation species generates more accurate ΔΔG estimations for T315A (Table S2), correctly predicting the sign of ΔΔG for both species of imatinib and the bulk estimated value. Despite Dasatinib only having a single protonation state, prospective methods obtain much more accurate ΔΔG values for L298F than for T315A (Table S3). The inaccuracy of ΔΔG predictions in the T315A predictions can be explained by an improper starting configuration with a trapped water molecule adjacent to residue 315, as described above (Fig. 7, Fig. S7–S9). Previous work has shown that considering the protonation state of titratable residues can improve sources of error in ΔΔG estimations.^101,102^ In contrast, we find that changing the protonation state of titratable amino acids does not improve ΔΔG accuracy relative to experiment (Table S5). However, previous work has highlighted the importance of protonation states in titratable residues in ABL1A kinase,^47,79,101^ indicating that while protonation state may be important for kinase activity, it may have less of an impact on direct inhibitor binding in the context of these two inhibitors. Overall, our work in light of previous findings highlights the importance of considering the protonation state of both titratable amino acids as well as of the ligand when predicting the impact of mutations.

Overall, using Perses to retrospectively analyze emblematic mutations that are challenging to estimate, we show that there are multiple potential sources in the starting configuration of a system that can introduce large systematic errors in estimations of mutation ΔΔG. These starting configuration errors can arise mainly in two ways: protonation state and through under-sampling of water interaction networks. Protonation states of both ligand and titratable amino acids must be considered when comparing with experiment, as it is important to consider both sources of variation in protonation states. This includes considering both tautomers as well as altered charge species of ligands and amino acids. We also show that appropriately sampling water networks is critical, as alchemical transformations can perturb these water networks without giving them time to appropriately relax and equilibrate. This can in turn introduce large errors in ΔΔG estimation. This can be remedied by longer sampling times, larger sampling windows, and ensuring appropriate time to relax between non-equilibrium alchemical transformation steps.

## CONCLUSIONS

In this work we highlight the ability of computational physics-based methods to prospectively estimate the thermodynamic cost of mutating an amino acid to predict the ΔΔG of ligand binding upon mutation. Using two different inhibitors of Abl kinase, imatinib and dasatinib, we show that NanoBRET measurements provide a consistent reproducible measurement for experimentally measuring ΔΔG for multiple constructs without having to rely on pooling measurements from a variety of sources and accruing multiple points of bias and error. This demonstrates that NanoBRET is a suitable tool to provide benchmarks for computational predictions of mutation-driven impact, and that this dataset will provide an initial test bed for the benchmarking of future predictive methods (SI data). We evaluate multiple prospective physics- and structure-based methods, alchemical free energy methods sampling via both replica exchange and nonequilibrium switching, Rosetta’s flex_ddg protocol, and a Random Forest model trained entirely on prior data. We show that all methods provide reasonably similar prediction accuracy, but that physics-based simulation provides improved minimum prediction accuracy on a per-residue level. Importantly, we emphasize that all structure-based and physics-based methods can reasonably propose whether or not a mutation is significantly resistant or sensitizing with an average accuracy of 0.81. Within this range of accuracy, these methods can act as suitable classifiers to prospectively predict whether a mutation is resistant or sensitizing in the absence of prior data. We also show that physics-based simulations are best able to capture the impact of distal mutations on ΔΔG of inhibitor binding, presumably due to their ability to capture the impacts of mutation on the dynamics of the kinase. Physics-based simulations are also more able to capture long-range electrostatic effects at greater distances, some approaches such as Rosetta and Random Forest models do not consider due to default cutoffs in their implementations. Lastly, we consider emblematically difficult mutations to predict and show that considering either 1) alternative starting configurations to better sample water-interaction networks, or 2) multiple protonation states and charge species, can improve ΔΔG prediction accuracy. By sharing these datasets, both experimental and computational predictions, in an open-source manner, we provide an initial test benchmark for future computational methods to evaluate their ability to predict the impact of resistant and sensitizing mutations. We hope these datasets highlight the utility of these methods and their capacity to classify the mechanistic impact of mutations in data-poor regimes where prior biochemical data is absent.

## ACKNOWLEDGEMENTS AND FUNDING

We want to thank the citizen-scientists of Folding@home for donating their computing resources for the Perses Free Energy estimations (projects 17606–17630). This work used resources from the High-Performance Computing Group at Memorial Sloan Kettering Cancer Center. The authors are grateful to the MSKCC DigITs and HPC team, especially Jamie Cheong, Lohit Valleru, and Monica Chakradeo for their assistance with high-performance computing resources. We are grateful to Ivy Zhang for helpful discussions regarding data visualization and feedback. SS is a Damon Runyon Quantitative Biology Fellow from the Damon Runyon Cancer Research Foundation (DRQ-14-22) and acknowledges support from a NCI Pathway to Independence Award for Outstanding Early-Stage Postdoctoral Researchers (NCI K99 CA286801). MA was supported by a Research Fellowship of the Alexander von Humboldt Foundation. JDC acknowledges funding from the National Institutes of Health (R35GM152017 and P30CA008748). DS acknowledges financial support from Bayer AG. AMR acknowledges funding from the National Institutes of Health (F30 CA260771). JDC, DS, and AV acknowledge funding from the Stiftung Charité and the BIH Einstein Foundation. MAS acknowledges funding from the National Institutes of Health (R35GM119437).

## DISCLOSURES

CDC, JPB, MA are employees of Bayer AG. CDC and MA are shareholders of Bayer AG. DS is an employee of Nuvisan. VG is an employee of Janssen pharmaceuticals and may own equity. JDC is a current member of the Scientific Advisory Board of OpenEye Scientific Software and Founder and CEO of Achira, and has equity interests in Achira and PICO Therapeutics. As a scientific advisor or invited speaker, he has received consulting or speaking fees from Abbvie, Astex, Boehringer-Ingelheim, Blueprint Medicines, Celgene, Foresite Capital, Foresite Labs, Interline Therapeutics, MPM Capital, OpenEye Scientific, Schrödinger, and Ventus Therapeutics. The Chodera laboratory receives or has received funding from multiple sources, including the National Institutes of Health, the National Science Foundation, the Parker Institute for Cancer Immunotherapy, Relay Therapeutics, Entasis Therapeutics, Silicon Therapeutics, EMD Serono (Merck KGaA), AstraZeneca, Vir Biotechnology, Bayer, XtalPi, Interline Therapeutics, the Molecular Sciences Software Institute, the Starr Cancer Consortium, the Open Force Field Consortium, Cycle for Survival, a Louis V. Gerstner Young Investigator Award, and the Sloan Kettering Institute. A complete funding history for the Chodera lab can be found at http://choderalab.org/funding.

## DISCLAIMER

The content is solely the responsibility of the authors and does not necessarily represent the official views of the National Institutes of Health.

## DATA AVAILABILITY AND SUPPLEMENTARY DATA

All input files, data outputs, and relevant structural files for prospective predictions, reported NanoBRET data, analysis notebooks, and mutation counting are available on OSF (https://osf.io/s6ktq/). Input and analysis scripts, structures, and notebooks for the retrospective analysis via Perses are also available in the same OSF repository (https://osf.io/s6ktq/). The full volume of Perses files, containing simulation data, work values, and hybrid topology factories are available freely upon request and require 1.9 TB of storage space. As such, the files are available upon request without question but are not available online due to the lack of a suitable online storage repository. All methods described above are open-source except FEP+, which is not open-source, and Rosetta, which is free to non-commercial users. Source code is available for GROMACS (https://www.gromacs.org/), PMX (https://github.com/deGrootLab/pmx), Perses (https://github.com/choderalab/perses), OpenMM (https://openmm.org/), and OpenMMTools (https://github.com/choderalab/openmmtools).

## AUTHOR CONTRIBUTIONS

Conceptualization: JDC, MAS

Methodology: JDC, MAS

Investigation: CDC, SS, DS, VG, MA, JB, JG, WG, SH

Writing -- Original Draft: SS, JB, JS,

Writing -- Review & Editing: SS, CDC, VG, DS, AV

Funding Acquisition: JDC, MAS, AV

Resources: JDC, MAS, BdG

Supervision: BdG, CDC, JDC, MAS, AV

## Notes

https://osf.io/s6ktq/

### Competing Interest Statement

The authors have declared no competing interest.

### Summary of Updates

1. Updated supplemental information and figures. 2. Updates to main text figures for improved clarity. 3. Updated main text for improved flow and readability

