## Supplemental-figures-information for "Prospective evaluation of structure-based simulations reveal their ability to predict the impact of kinase mutations on inhibitor binding"

**AFFILIATIONS:**
[1] Computational and Systems Biology Program, Memorial Sloan Kettering Cancer Center, New York, NY, 10065, USA
[2] Computational Chemistry, Janssen Research & Development, Turnhoutseweg 30, Beerse 2340, Belgium
[3] Computational Biomolecular Dynamics Group, Department of Theoretical and Computational Biophysics, Max Planck Institute for multidisciplinary sciences, D-37077 Göttingen, Germany
[4] In Silico Toxicology and Structural Bioinformatics, Institute of Physiology, Charité – Universitätsmedizin Berlin, corporate member of Freie Universität Berlin and Humboldt-Universität zu Berlin, Augustenburger Platz 1, 13353 Berlin, Germany
[5] Department of Pharmacological Sciences, Stony Brook University Medical School, Stony Brook, NY 11794, United States
[6] Tri-Institutional PhD Program in Computational Biology and Medicine, Weill Cornell Graduate School of Medical Sciences, Cornell University, New York, NY 10065, United States
[7] Structural Biology & Computational Design, Research and Development, Pharmaceuticals, Bayer AG, 13342 Berlin, Germany
[8] Open Molecular Software Foundation, Davis, CA 95618, USA
[9] Department of medicine II, University Hospital Aachen, Pauwelsstraße 30, 52074 Aachen, Germany
[10] Data Driven Drug Design, Faculty of Mathematics and Computer Sciences, Saarland University, 66123 Saarbrücken, Germany

§Present address: Nuvisan ICB, Life Science Chemistry, Digital Müllerstraße 178, 13353 Berlin, Germany
^Present address: Computational Chemistry, Janssen Research & Development, Janssen Pharmaceutica N. V., Turnhoutseweg 30, B-2340 Beerse, Belgium
†Present address: Bayer Research and Innovation Center, 238 Main St, Cambridge, MA, 02142, USA.

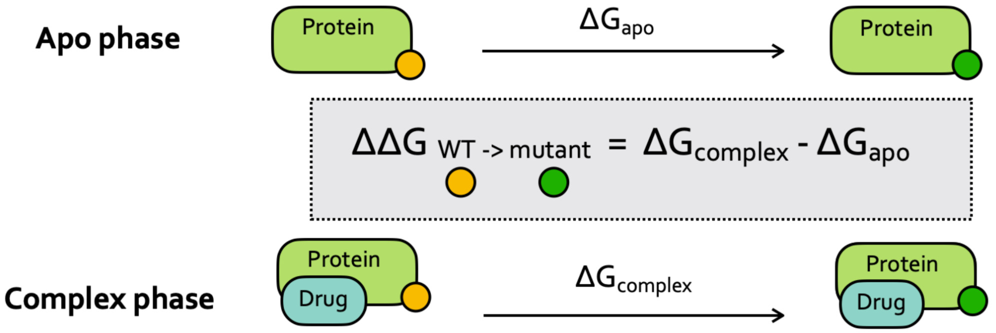


**Figure S1.** Representative thermodynamic cycle that highlights the set of transformations used to predict the ΔΔG of mutations upon drug binding in a protein-ligand system.[47] Transformations are computed by computing the energetic cost of mutating a residue from wild type (yellow) to mutant (dark green), in both apo phase (top row) and in complex with drug binding (bottom row). From these calculations, ΔΔG can be computed by subtracting the apo phase ($\Delta G_{apo}$) from the complex phase ($\Delta G_{complex}$).

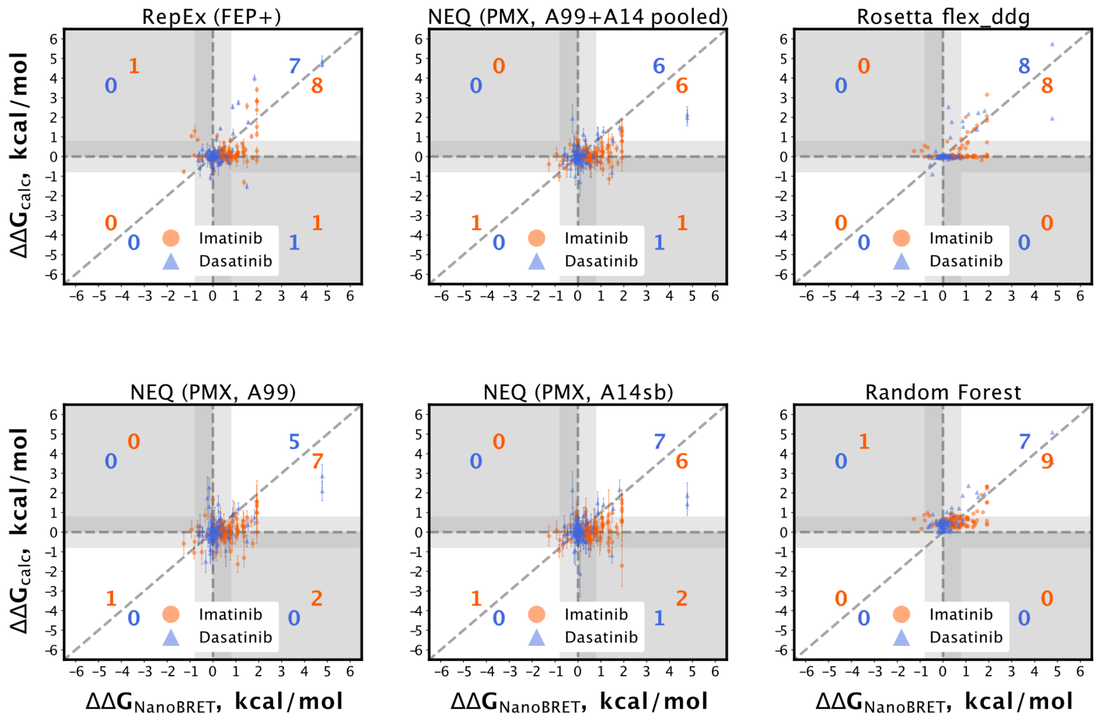


**Figure S2. Truth tables for each method demonstrate the capacity of computational methods to act as classifiers.** The number of "true positives" of ΔΔG predictions for a variety of methods are shown (top right and bottom left integers). A prediction is considered a true positive if its has the same sign as the experimental NanoBRET predictions, and both have |ΔΔG| > 1 kcal/mol (top right and bottom left quadrants). True positives for resistant (top right integers) and sensitizing mutations (bottom left integers) are shown for both imatinib (orange) and dasatinib (blue). Experimental ΔΔG measurements with magnitude below 1 kcal/mol are not labeled as "true positives." Prospective methods shown are Replica Exchange using FEP+ (top left), Nonequilibrium switching using PMX using Amber99 force field (bottom left), the Amber14sb force field (bottom middle), and the resultant prediction taken from pooling the work values from both force fields (top middle). Rosetta's flex_ddg (top right) and a random forest model trained on prior data (bottom right) are also shown.

**Table S1.** Summary statistics from AUPRC curves highlighting the similarity in performance for each method. The 95% confidence intervals are calculated based on bootstrapping 1000 repeats with replacement

| Method | Pooled AUPRC | 95% Conf. Interval | Pooled accuracy | Distance from imatinib baseline at 1 kcal/mol | Distance from dasatinib baseline at 1 kcal/mol |
|---|---|---|---|---|---|
| **FEP+** | 0.6 | 0.45–0.73 | 0.82 | 0.27 | 0.68 |
| **PMX A99/A14** | 0.55 | 0.38–0.71 | 0.82 | 0.52 | 0.57 |
| **PMX A99** | 0.47 | 0.31–0.65 | 0.81 | 0.66 | 0.63 |
| **PMX A14** | 0.52 | 0.37–0.68 | 0.82 | 0.36 | 0.46 |
| **Rosetta 15** | 0.6 | 0.44–0.77 | 0.85 | 0.41 | 0.54 |
| **Random forest (ML)** | 0.58 | 0.44–0.71 | 0.79 | 0.19 | 0.44 |

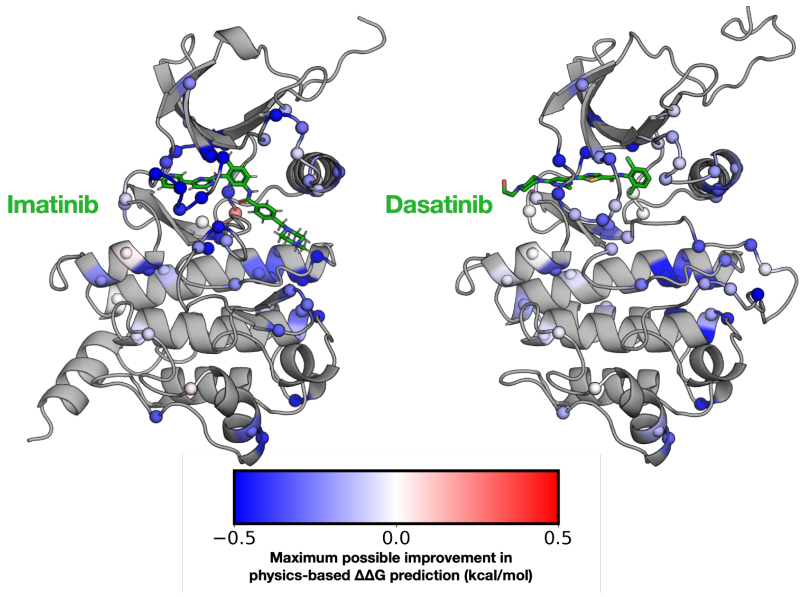


**Figure S3. Maximum possible improvement in ΔΔG prediction by an alchemical simulation method relative to a non-alchemical method.** The maximum possible improvement the is computed by taking the least accurate non-alchemical ΔΔG prediction and subtracting it from the most accurate ΔΔG prediction, where a negative score indicates the best possible improvement for any predicted ΔΔG value. These values are mapped onto the structure of Abl kinase (PDB: 1OPJ) for each mutation (spheres), and colored to indicate the degree to which alchemical methods are able to improve upon individual non-alchemical predictions (color scale, below). Mappings are made for both imatinib predictions (left) and dasatinib (right).

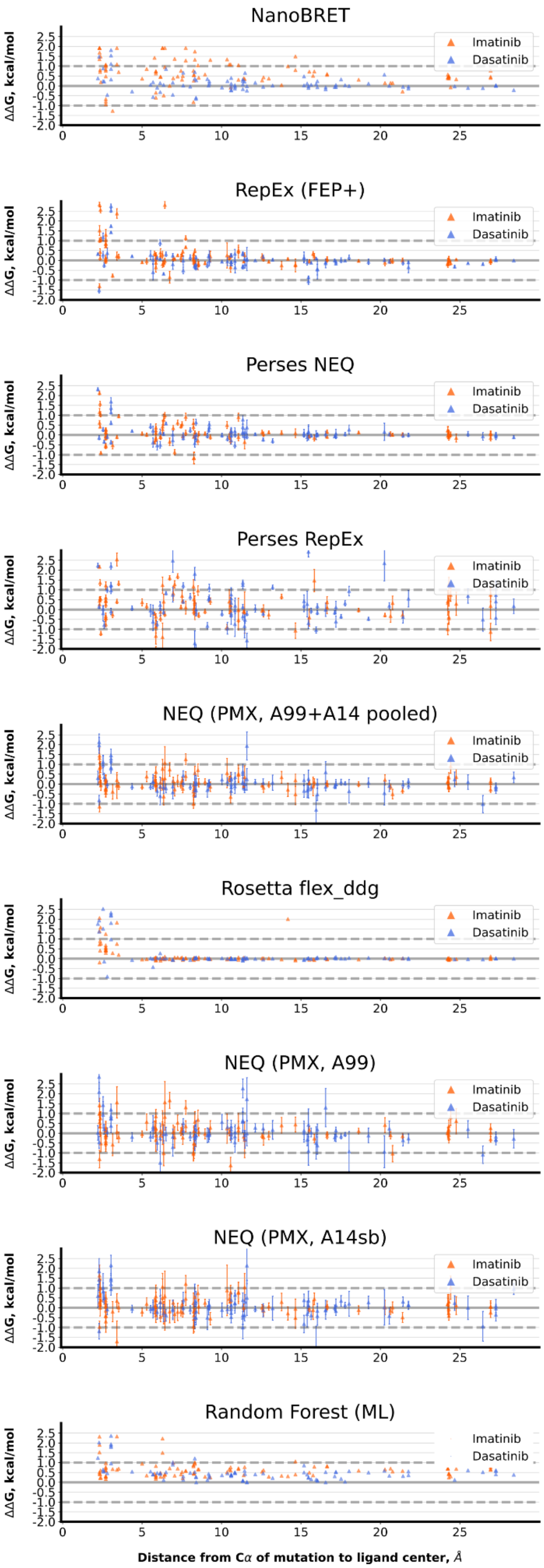


**Figure S4. ΔΔG estimations from both experiment and computation.** Computed ΔΔG for NanoBRET and each computational method is plotted for imatinib (orange) and dasatinib (blue) as a function of the distance from the residue's Cα carbon to the center of mass of the ligand in the crystal structure.

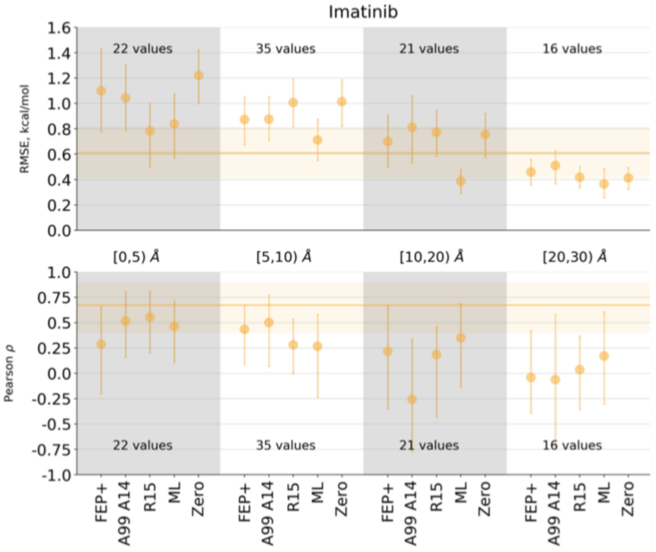


**Figure S5**. **Estimation accuracy and summary statistics is dependent on the degree of distance from the active site.** Calculation accuracy grouped in ranges of the distance between the mutated residue and the imatinib. “Zero” denotes a prediction where every mutation is predicted to be neutral in impact on inhibitor binding (ΔΔG = 0 kcal/mol). Horizontal lines mark RMSE and correlation values between two experimental measurements.

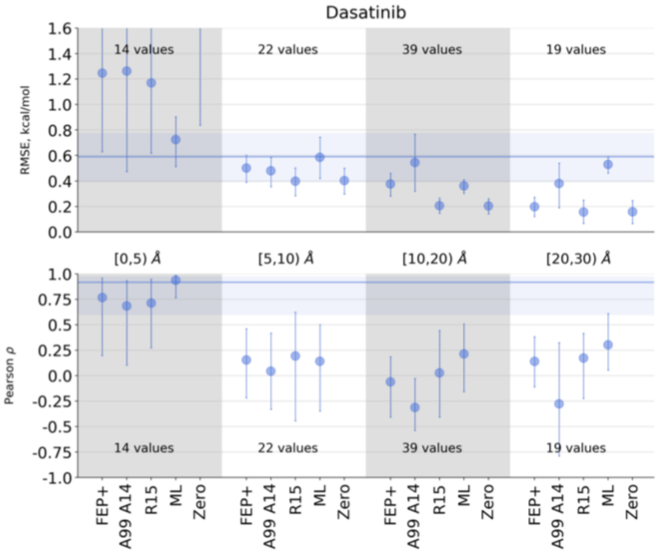


**Figure S6**: Calculation accuracy grouped in ranges of the distance between the mutated residue and the inhibitor. “Zero” denotes a prediction where every mutation is predicted to be neutral in impact on inhibitor binding (ΔΔG = 0 kcal/mol). Horizontal lines mark RMSE and correlation values between two experimental measurements (Fig. 2).

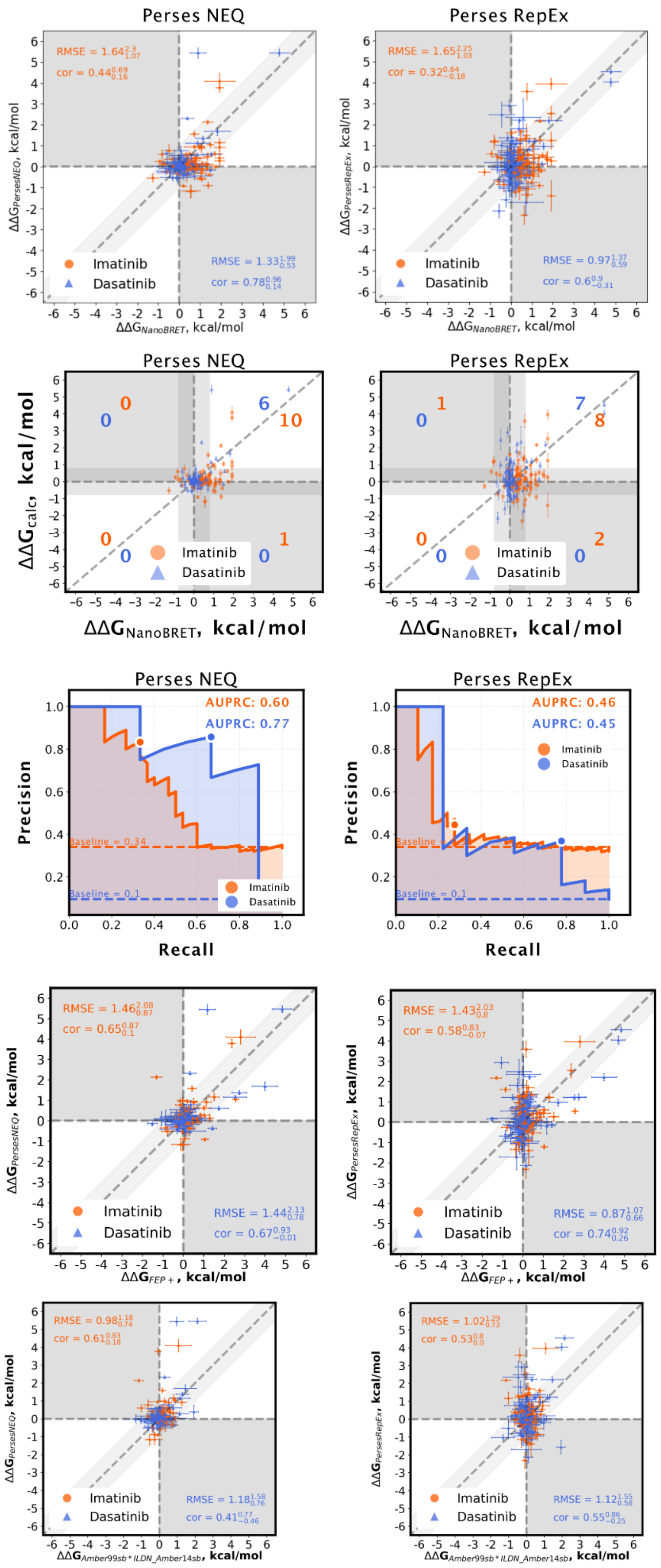


**Figure S7. Open-source tools like Perses and PMX can estimate ΔΔG and act as a classifier while allowing a user to robustly tune parameters and investigate simulation details.** Scatterplots (top row) showing predictions made using Perses using both NEQ (left column) and Replica Exchange (RepEx, right column), including both Root Mean Square Error (RMSE) and Pearson correlation (labeled "cor"). These can be converted into truth tables (second row) that generate Precision-Recall curves (third row) for both sampling methods on Perses are also shown for both Imatinib (orange) and dasatinib (blue). These Perses-based calculations using both NEQ and Replica Exchange sampling are compared against other physics-based methods FEP+ (fourth row) and PMX (A99 + A14 pooled) (bottom row) to evaluate consistency of predictions across retrospective and prospective methods with respect to RMSE and Pearson correlation.

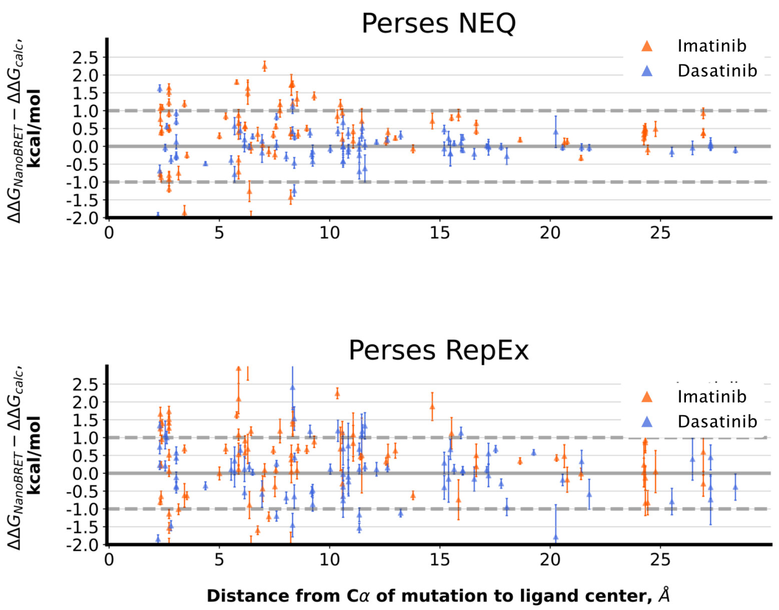


**Figure S8. Perses-based alchemical simulations ΔΔG estimations are also able to predict the impact of distal mutations on imatinib and dasatinib binding.** The deviation from predicted ΔΔG to experiment is plotted for imatinib (orange) and dasatinib (blue) as a function of the distance from the residue's Cα carbon to the center of mass of the ligand in the crystal structure. Values are shown for predictions made with Perses using Nonequilibrium Cycling (top) and replica exchange (bottom).

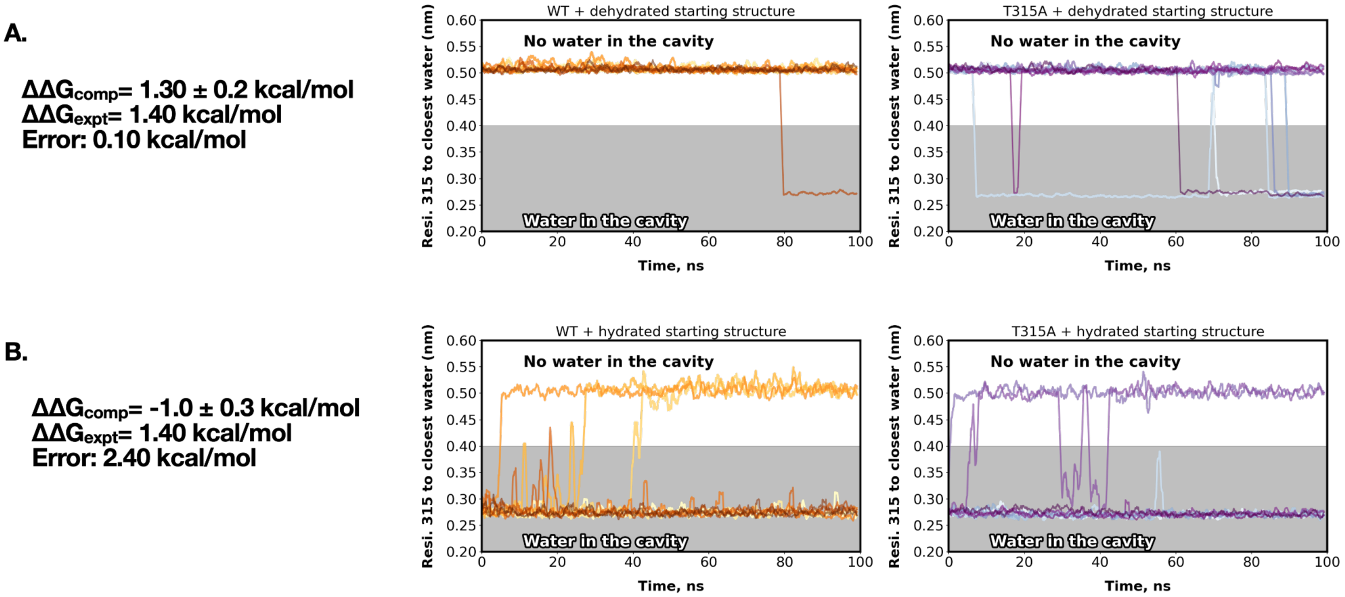


**Figure S9. Trajectories started with the crystal water removed both increase alchemical ΔΔG prediction and decreased water binding. A.** The distance from residue 315 to the closest water molecule in equilibrium MD trajectories (right) started from a crystal structure that removed the closest water molecule to residue 315, rendering the residue in a "dehydrated" state. Trajectories were collected for both wild-type (middle) and T315A constructs (right) Alchemical free energy calculations estimating ΔΔG using this structure yield accurate predictions (left). **B.** Distance from residue 315 to the closest water molecule in equilibrium trajectories starting from the same crystal structure with all water molecules present (middle and right) show increased proximity and potential interaction between residue 315 and water molecule in both WT (middle) and T315A constructs (right). However, alchemical ΔΔG predictions made using this starting structure are far less accurate (left).

**Table S2. Protonation state of imatinib alters ΔΔG predictions for several simulated mutations.** Predictions were considered significantly changed when the error in predicted ΔΔG compared to the simulations with positively charged imatinib changed by 0.6 kcal/mol or more. Reported values of ΔΔG calculations are the mean of at least 3 independent repeats. Each cell is colored by how much the prediction deviates from the NanoBRET (bottom row of table).

| **Mutation** | Ligand and net charge | NanoBRET (kcal/mol) | FEP+ (kcal/mol) | a99 | a14 | PersesNEQ | PersesRepEx |
|---|---|---|---|---|---|---|---|
| **L298F** | imatinib+0 | **-1.27** | -0.77 | -0.58 | -0.2 | -0.55 | 0.28 |
| **L298F** | imatinib+1 | | | | | -0.79 | -2.91 |
| **L298F** | Imatinib (bulk) | | | | | -0.77 | -2.86 |
| **T315A** | imatinib+0 | **1.36** | -1.31 | -1.31 | -0.97 | 2.01 | 2.48 |
| **T315A** | imatinib+1 | | | | | 2.36 | 2.45 |
| **T315A** | Imatinib (bulk) | | | | | 2.30 | 2.45 |
| | | Colorscale for difference between computational prediction vs. NanoBRET measurement: (kcal/mol) | < 0.5 | 0.5 - 1.0 | 1.0 - 1.5 | > 1.5 | |

**Table S3. Dasatinib alters ΔΔG predictions for several simulated mutations.** Predictions were considered significantly changed when the error in predicted ΔΔG compared to the simulations with positively charged imatinib changed by 0.6 kcal/mol or more. Reported values of ΔΔG calculations are the mean of at least 3 independent repeats. Each cell is colored by how much the prediction deviates from the NanoBRET (bottom row of table).

| **Mutation** | Ligand | NanoBRET (kcal/mol) | FEP+ (kcal/mol) | A99 (kcal/mol) | A14 (kcal/mol) | PersesNEQ (kcal/mol) | PersesRepEx (kcal/mol) |
|---|---|---|---|---|---|---|---|
| **L298F** | dasatinib | **-0.31** | 0.07 | -1.5 | 0.27 | -0.47 | 0.60 |
| **T315A** | dasatinib | **1.48** | -1.51 | -0.46 | -1.19 | 2.39 | 2.24 |
| | | Colorscale for difference between computational prediction vs. NanoBRET measurement: | < 0.5 | 0.5 - 1.0 | 1.0 - 1.5 | > 1.5 | |

**Table S4. A neutral protonation state of imatinib alters FEP+ predictions for several simulated mutations**. Reported values of FEP+ calculations are the mean of at least 3 independent repeats.

| | | **FEP+ prediction: ΔΔG ± SD (kcal/mol)** | | |
|---|---|---|---|---|
| **Mutation:** | **Experiment value (kcal/mol)** | **Prediction for imatinib$^{+1}$** | **Prediction for imatinib$^{+0}$** | **Bulk Imatinib prediction** |
| **Y353H** | 1.367 | 0.077 ± 0.059 | 0.880 ± 0.220 | 0.113 ± 0.070 |
| **F359I** | 1.068 | 0.033 ± 0.284 | 0.733 ± 0.317 | 0.066 ± 0.286 |
| **N368S** | 0.002 | -0.853 ± 0.522 | -0.180 ± 1.146 | -0.825 ± 0.553 |
| **E282G** | 1.284 | 0.673 ± 0.090 | 0.007 ± 0.352 | 0.582 ± 0.107 |
| **E282K** | 1.920 | 1.130 ± 0.198 | 0.173 ± 0.345 | 0.966 ± 0.208 |
| **V289F** | 0.710 | 0.430 ± 0.270 | 2.02 ± 0.435 | 0.475 ± 0.281 |
| **E292V** | 0.388 | 0.080 ± 0.227 | -0.540 ± 0.220 | -0.001 ± 0.225 |
| **M351K** | 1.920 | 1.920 ± 0.303 | 0.440 ± 0.422 | 1.546 ± 0.311 |
| **E355G** | 0.539 | -0.017 ± 0.330 | -0.740 ± 0.169 | -0.120 ± 0.315 |

**Table S5. Alternative side chain protonation states do not improve FEP+ predictions.** Predictions were considered significantly changed when the error in predicted DDG compared to the simulations with the default side chain protonation state changed by 0.6 kcal/mol or more. Protonation state of imatinib was +1. Reported values of FEP+ calculations are the mean of at least 3 independent repeats.

| | | Experimental | Default protonation state | | Alternative protonation state | |
|---|---|---|---|---|---|---|
| **Mutation** | **Ligand** | **ΔΔG [kcal/mol]** | **Mutation** | **ΔΔG [kcal/mol] ± SD** | **Mutation** | **ΔΔG [kcal/mol] ± SD** |
| Y253H | imatinib | 1.920 | TYR253HID | 1.517 ± 0.429 | TYR253HIE | 0.877 ± 0.452 |
| E282K | imatinib | 1.920 | GLU282LYS | 1.130 ± 0.198 | GLU282LYN | 0.260 ± 0.110 |
| M351K | imatinib | 1.920 | MET351LYS | 1.920 ± 0.303 | MET351LYN | 0.017 ± 0.442 |
| L248R | dasatinib | 0.883 | LEU248ARG | 1.184 ± 0.468 | LEU248ARN | 2.230 ± 0.184 |
| G250E | dasatinib | 0.187 | GLY250GLU | 0.028 ± 0.416 | GLY250GLH | -0.877 ± 0.685 |

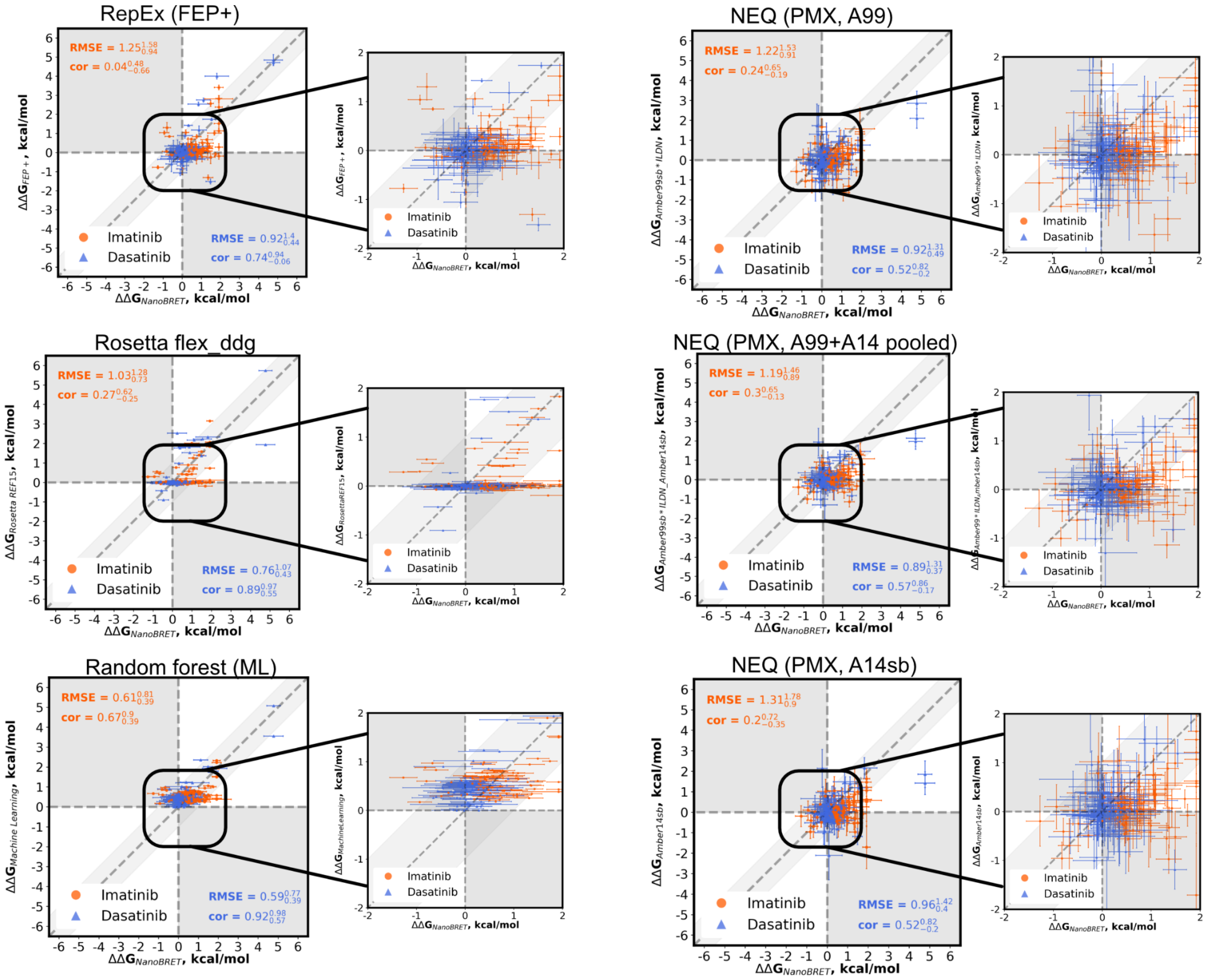


**Figure S10. Prospective computational methods have many predictions centered around zero.** Scatterplots showing the prospective ability of different computational methods and their ability to predict the ΔΔG of either imatinib (orange) or dasatinib (blue) binding. Inset plots for each scatterplot (ranging from -2 to +2 kcal/mol), are also shown adjacent to the relevant scatter plot (black inset outline). These comparisons are all done relative to the same ΔΔG measurements collected using NanoBRET (x-axis). Prospective methods shown in the left column are Replica Exchange using FEP+ (top row), Rosetta's flex_ddg (middle row) and a random forest model trained on prior data (bottom row). In the right column, prospective methods shown are Nonequilibrium switching using PMX using the Amber99 force field (top row), the Amber14sb force field (bottom row), and the resultant prediction taken from pooling the work values from both force fields (middle row). are also shown. Root Mean Square Error (RMSE) and Pearson correlation (labeled "cor") are provided in the top left (imatinib) and bottom right corners (dasatinib).
